# Machine learning techniques for classifying the mutagenic origins of point mutations

**DOI:** 10.1101/342618

**Authors:** Yicheng Zhu, Cheng Soon Ong, Gavin A Huttley

## Abstract

There is increasing interest in developing diagnostics that discriminate individual mutagenic mechanisms in a range of applications that include identifying population specific mutagenesis and resolving distinct mutation signatures in cancer samples. Analyses for these applications assume that mutagenic mechanisms have a unique relationship with neighboring bases that allows them to be distinguished. Direct support for this assumption is limited to a small number of simple cases, e.g. CpG hypermutability. We have directly evaluated whether the mechanistic origin of a point mutation can be resolved using only sequence context for a more complicated case. We contrasted mutations originating from the multitude of mutagenic processes that normally operate in the mouse germline with those induced by the potent mutagen N-ethyl-N-nitrosourea (ENU). The considerable overlap in the mutation spectra of these two samples make this a challenging problem. Employing a new, robust log-linear modeling method, we demonstrate that neighboring bases contain information regarding point mutation direction that differs between the ENU-induced and spontaneous mutation classes. A logistic regression classifier proved to be substantially more powerful at discriminating between the different mutation classes than alternatives. Concordance between the feature set of the best classifier and information content analyses suggest our results can be generalized to other mutation classification problems. We conclude that machine learning can be used to build a practical classification tool to identify the mutation mechanism for individual genetic variants. Software implementing our approach is freely available under the BSD 3-clause license.

## Introduction

In most catalogs of genetic variation, the data consist of variants that derive from a mixture of mutagenic processes. Whether analysis of the genetic variants alone allows resolving the causative mechanism for an individual genetic variant remains an open question. Instances of a singular etiological relationship between point mutation mechanism and flanking sequence are known for only a small number of relatively simple cases. From a biochemical perspective, it seems a reasonable conjecture that the sequence of neighboring bases should affect mutagenic processes in general. This conjecture remains substantively unverified as is the related conjecture that knowledge of neighboring sequence is sufficient to identify the specific mutagenic origin. Methods have been developed that can discriminate between entire mutation spectra (Zhu et al., 2017), such as those characteristic of cancers, and to estimate the major components of these spectra (Alexandrov et al., 2013; Shiraishi et al., 2015). As far as we are aware there has not been a detailed examination of the relationship between a mutation mechanism and neighboring bases with a view to identifying mechanistic origins of individual variants. Here, we employ machine learning methods to address this using a data set of point mutations of known origin. We limit discussion, and analysis, to the 12 distinct single nucleotide point mutations.

In mammals, mutation processes exhibit considerable heterogeneity which manifests between genomic locations, cell types, disease states and clinical treatments. The within-genome heterogeneity of sequence composition is taken as an indicator of the heterogeneous operation of mutation processes operating in the germline and multiple factors are implicated in driving this pattern (for review see Hodgkinson and Eyre-Walker, 2011). These include factors that distinguish gametogenesis between the sexes (e.g. Huttley et al., 2000), which manifest at the level of entire chromosomes, to the localized operation of transcription-coupled DNA repair processes (e.g. Svejstrup, 2002). There is also considerable complexity in the origin of mutations affecting somatic tissues. Variation in mutagenesis distinguishes normal cell lineages, as evidenced by the biochemically specified somatic hypermutation that occurs in immune cells (Chahwan et al., 2012). The spectrum of mutations can be a distinctive feature of different cancers (Pleasance et al., 2010). This may result from tissue-specific exposure to exogenous mutagens, such as the reported excess of G → T* transversions (where * indicates a mutation direction and its strand complement) in smoking-associated lung cancer (Hainaut and Pfeifer, 2001). It may also reflect defects in specific DNA repair processes (Viel et al., 2017). In all of these cases the catalog of mutations arises from a mixture of different processes, making assignment of a specific cause to a single mutation challenging.

Germline heterogeneity in mutagenesis has been correlated with a number of genomic features and processes including the abundance of G and C nucleotides (hereafter GC) and sexual dimorphism in gametogenesis. The primary explanation for the positive correlation with GC is that it reflects a causal relationship with the recombination rate via the process of biased gene conversion (Hodgkinson and Eyre-Walker, 2011; Meunier and Duret, 2004; Hellmann et al., 2005). Differences between the sexes in the spectrum of point mutations leads to differences in GC between chromosomes based on time spent in the male germline (Huttley et al., 2000).

We can decompose the process of a mutation into two fundamental steps: lesion formation followed by a failure of DNA repair to reconstitute the original base pair. High exposure of cells to UV light, which elevates formation of dipyrimidine lesions, illustrates the role of lesion creation on mutagenesis (Pfeifer et al., 2005). The accumulation of defects in DNA mismatch repair genes, which contribute to development of colorectal cancer, illustrate the role of defective DNA repair (Viel et al., 2017). In both of these cases, the rate at which the different point mutations occur can be affected, highlighting that different types of point mutation can have a common mechanistic origin. As systemic changes to mutation process are a feature of cancer cells, a primary analysis focus in cancer biology has been to resolve mutagenic signatures that characterize cancers (Alexandrov et al., 2013; Shiraishi et al., 2015). This work exploits the presumed relationship between point mutation processes and flanking DNA sequence.

The nucleotides flanking a mutated position contain information regarding the mutagenesis process responsible for the change. Hypermutability of the CpG dinucleotide illustrates the relationship between neighboring bases and point mutation mechanism. Association of a 3’-G with elevated C→T mutation rates derives from the binding preference of DNA methylases (Krawczak et al., 1998). These enzymes bind to this dinucleotide and modify C to 5-methyl-cytosine. The resulting modified base exhibits a 10-fold increase in spontaneous deamination rate, an effect so pronounced as to almost entirely swamp alternate causes of C→T mutations (Zhu et al., 2017). The apparent simplicity of the relationship between C→T point mutations and flanking nucleotides reflects the dominance of a single chemical process in creating lesions.

The sequence motifs associated with non-C→T point mutations are more complicated (Zhu et al., 2017), suggesting contributions from multiple mutagenesis mechanisms. It was shown from an analysis of millions of human germline mutations that more than one nucleotide at flanking positions were associated with the non-C→T point mutations (Zhu et al., 2017). This is consistent with multiple mutation mechanisms contributing to these point mutations. At present, the mechanistic basis underlying these mutation associated sequence motifs (mutation motifs) remains unknown. Even in the case of cancer, the diversity of defects in DNA repair that afflict these cells limit our certainty regarding the possible mechanisms that may be responsible for a specific genetic variant.

The systematic use of mutagens in forward genetic screens provides an opportunity to develop an understanding of the relationship between neighboring sequence and mutagenesis. N-ethyl-N-nitrosourea (ENU) is a synthetic alkylating chemical widely employed in mutagenesis studies (Álvarez et al., 2003; Lee et al., 2012; Stottmann and Beier, 2014), causing new germline mutations at ~100 times higher rate than the spontaneous mutation rate (Stottmann and Beier, 2014). Exposure to ENU can induce formation of a number of alkylation adducts including N^1^-adenine (e^1^A), O^4^-thymine (e^4^T), O^2^-thymine (e^2^T), and O^2^-cytosine (e^2^C) (Shrivastav et al., 2010; Noveroske et al., 2000). If the DNA repair system fails in repairing these adducts, they are mispaired during DNA replication to a non-complementary nucleotide, resulting in a single base change mutation (Noveroske et al., 2000; Justice et al., 1999). The resulting ENU-induced mutations are dominated by A → G* and A → T* mutations, with rare reported occurrences of C → G* mutations (Takahasi et al., 2007).

Whether ENU mutagenesis induces mutations randomly with regards to flanking DNA sequence is debated (Barbaric et al., 2007; Bauer et al., 2015). The unique ENU-induced mutation spectra distribution described above has provided the basis for the ENU-induced variant filtering strategy (Andrews et al., 2012). For example, removing any C → G* transversions, leaves only genetic variants likely to be generated by ENU process and thus candidates for novel phenotypes. We refer to this filtering strategy as the naïve (classification) method, in which the mutation mechanism is assigned solely on the basis of mutation direction. The approach has high accuracy solely because of the excess of ENU-induced mutations. However, there remains a possibility of misclassification of mutation origin in these studies as some fraction of the point mutations labeled as being ENU-induced will instead have originated by non-ENU mutagenesis. If sequence neighborhood does affect mechanism, then mutation classification techniques that exploit this information should improve over the naïve method.

Machine learning techniques are well suited to the problem of sequence-based classification of samples (James et al., 2013; Ben-Hur et al., 2008). The goal of machine learning classification is to find a rule, based on observed object features, that can assign new objects to one of several classes (James et al., 2013; Sonnenburg, 2008). Machine learning techniques have been applied to a diverse array of sequence-based classification problems ranging from microbial taxon assignment (e.g. Bokulich et al., 2018) to predicting the position of nucleosomes in eukaryotic cells from ChIP-seq data (e.g. Peckham et al., 2007).

In this study, we evaluate whether sequence features can improve the performance of classifiers devised to discriminate between mutagen induced and spontaneous point mutations in the mouse germline. We affirmed a highly significant influence of neighboring nucleotides on ENU point mutations and that these associations differ from those evident in spontaneous mutations. Our results reveal that a combination of *k*-mer size and representation of second-order interactions among nucleotides was able to markedly improve classification performance in comparison to the naïve classifier approach. All scripts developed for this work are made available under an open source license.

## Results

### Distinctions between ENU-induced and spontaneous point mutations

A logical requirement for using sequence features to discriminate samples is that those features differ in abundance between the samples. We addressed this using two complementary formal hypotheses tests. The “spectra” hypothesis test compares the distribution of point mutation outcomes in the two source materials. The “neighborhood” hypothesis test contrasts the association of neighboring bases with those point mutation outcomes. In both cases, ENU-induced germline point mutations were obtained from the Australian Phenomics Facility, and spontaneous germline mutations from Ensembl database (see Materials and Methods).

We employed a log-linear model to test the null of equivalence in mutation spectra between the ENU-induced and spontaneous samples (Zhu et al., 2017). This test considers the relative distribution of outcomes from mutations of, for example, the base T. A separate test was employed for each possible starting base. Consistent with published reports, the spectra of ENU-induced and spontaneous point mutations in the mouse were significantly different (Fig S1 and Table S1). To simplify the following, and as stated in the Introduction, we abbreviate the description of a point mutation and its strand complement using the notation X→Y*, i.e. A→G* refers to both A→G and its strand complement T→C. Direct examination of counts for the ENU-induced mutations reveals they were dominated by A → G* and A → T* mutations, with frequencies of 42% and 27% respectively. These contrast with their abundance in mouse spontaneous mutations of 29% and 3.7% respectively. Visualization of the spectrum analyses (Fig S1) reflects these changes in proportion. These differences affirm the basis for the current naïve mutation classification algorithms applied to ENU samples.

The striking difference in mutation spectra was also accompanied by striking differences in the magnitude and identity of neighboring base influences. Prior to discussing the results, we briefly describe the log-linear modeling analyses employed. We use position indices that are relative to the point mutation location, defined as position 0, with negative / positive indices representing 5’- / 3’- positions respectively. Consider, for example, the question of whether bases at the position immediately 3’- to a point mutation of A→G associate with the mutation. The test assesses the null hypothesis that in sequences where an A→G mutation occurred, the base counts at the +1 position are equivalent to those at the +1 position for occurrences of A in the reference distribution. This is an example of a single position (first-order), or independent position (denoted I in our modeling notation) effect. We can also evaluate whether the joint counts of bases at two positions are equal between the mutated and reference sequence collections (second-order dependence, or 2D). Our previous analyses of spontaneous germline mutations from humans identified neighbor effects as highly influential, and that independent and second-order effects dominated higher-order effects (Zhu et al., 2017). These analyses are readily extended to comparing equivalence between samples, as is the objective here. See Materials and Methods for more details.

Our analyses established there were strongly significant differences between the ENU-induced and spontaneous mutations in the identity of the associated mutation motifs, and their relative magnitude. To simplify the exposition, we limit our discussion here to description of the results from the A→G* case, the most abundant ENU-induced point mutation. (We note that all point mutations exhibited strongly significant differences and summarize these in Table S2). The maximum relative entropy (RE) association of independent positions with A→G was 5-fold larger in the ENU-induced sample. This maximum association was at +1 in the ENU-induced sample, compared with −1 for the spontaneous sample (Fig S1). Using the log-linear model, we rejected the null hypothesis of the equivalence between ENU-induced and spontaneous samples for neighboring base associations with A→G mutations. While these samples revealed highly significant differences for nearly all effects orders (Table S1), the magnitude of difference was greatest for the I and 2D effects (Fig S2). As mentioned above, these patterns held true for all point mutation directions (Table S2).

Of further relevance to feature selection for classifier design is the physical limit to these associations. Estimation of the physical limit of association from longer flanking contexts was obtained using relative entropy as per Zhu et al. (2017) (see Fig S3 and Table S3). The ENU-induced sample showed the physical limit mean, median and standard deviation of 3.2bp, 2bp, and 1.7bp respectively. In contrast, the corresponding statistics for spontaneous mutations were 2.9bp, 2.5bp, and 2bp. As a consequence of this variability, we considered a range of different neighborhood sizes in development of the classifiers.

### Development of a two-class machine learning classifier

In developing classifiers, we evaluated a collection of algorithms, sample sizes, sequence feature sets, *k*-mer size and hyperparameter values (see Materials and Methods for details). Classifier development was strictly limited to data from a single mouse chromosome. We arbitrarily chose chromosome 1 given availability of sufficient data (see Table S4). We note here that we present only the Logistic Regression (LR) classifier results in the manuscript. LR was chosen because of its systematically better performance than the naïve Bayes (NB) classifiers and interpretability of the resultant classifiers, compared to XGBoost. It is noteworthy that XGBoost exhibited a superior learning response compared to LR. When applied to the genome, however, the advantage of XGBoost over LR was weak. This drop in performance likely arises from substantial over-fitting by XGBoost. (See Fig S4 for the learning curves from the best classifiers for each algorithm.)

Unless indicated otherwise, classifier performance was measured as the area under the receiver operating characteristic curve (AUC) score (see Material and Methods for more detailed justification of this choice). For any particular classifier, its performance was measured using the mean and standard error derived from 10 replicate AUC measures obtained from the cross validation analysis. A classifier whose mean AUC score was greater than that of another classifier was taken to be superior, after considering the standard errors.

In the following, we describe the classifier feature sets using a combination of the terms M, I, 2D, 2Dp, FS and GC%. We refer the reader to Materials and Methods for a more detailed description. These terms correspond to the mutation direction (M), the set of contributions from independent flanking positions (I), and the set of contributions arising from 2-way dependence among flanking positions (2D). The 2Dp notation refers to a subset of 2D where the positions are physically proximal to each other and/or the mutating site. The fully saturated (FS) model is a model containing M and all possible independent and multi-position interactions. (In the regressions, the exact values for the I and D terms depend on the value of *k*.) The GC% corresponds to the percentage of G+C nucleotides in flanking DNA sequence.

For LR, we made choices regarding two hyperparameters. *ℓ*_1_ regularization was chosen as it prunes out unneeded features by setting their associated weights to 0 (Bühlmann and Van De Geer, 2011). This allowed us to establish which features contribute to the classification. The regularization parameter *C* controls overfitting by affecting the trade-off between variance and bias of regression parameter estimates. We selected the value of *C* that returned the best classifier performance on the validation set (see Materials and Methods).

Comparison of training curves resulting from classifier evaluation indicated that M+I+2D provided robust performance. The learning curves show the sensitivity of the classifier performance to training set size, where the latter is the total of both ENU-induced and Spontaneous classes. For the categorical feature sets, we considered four distinct models: M, M+I, M+I+2D and FS. It can be seen from Fig S2 that when training size is > 4, 000 samples, the rate of classifier performance improvement with increasing sample size dropped off markedly. For subsequent comparisons, we used classifiers trained on data sets with ~16,000 samples as their standard errors allowed greater resolution between the feature sets. Of the classifiers that only included categorical features, the naïve classifier employed for classifying ENU-induced mutations, M, was the least accurate. Inclusion of individual position features, represented by I, provided a substantial improvement over M. The best performing classifiers, however, included features representing dependence among positions (see Table S5 for detailed statistics). That said, the overlap in standard errors of the 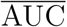 for the M+I+2D and FS models (Fig S2) indicate that inclusion of two-way dependence captured the majority of information contained by the sequence neighborhood. The value of *C* that returned maximal performance was consistently 0.1 for all models and all samples that considered higher-order interactions (i.e. 2D and above).

### Choosing neighborhood size

As illustrated by the log-linear analyses reported above, the physical limit of neighboring base influence differs between point mutations and mutation mechanism (Figure S3 and Table S2). Recalling that a symmetric neighborhood size of 3 equates to *k* = 7, we initially assessed the impact of sequence neighborhood size by comparing performance for three different *k*-mer sizes (3, 5, 7) for the M+I and M+I+2D feature sets. Comparison of learning curves established that for training set sizes > 4, 000, classifiers based on a 7-mer context performed better than the other two values of *k* (Fig S3). The impact of choice of *k* differed between the feature sets, with the strongest improvements with increasing *k* evident for the M+I+2D model. These results suggested exploration of larger *k*. Initial efforts at modest *k* failed due to excessive memory requirements as the number of 2D parameters increases *k*^2^. The log-linear results presented here and previously (Zhu et al., 2017), indicated that most information arising from interactions between positions is captured by just proximal positions. Accordingly, we considered the 2Dp feature subset (see Materials and Methods) where an interaction between two positions was included only if they were physically adjacent to each other, or straddled the mutating base. For analyses with *k* > 7, we considered just M+I and M+I+2Dp feature sets. The results reinforced our choice of the M+I+2Dp feature set and identified *k* = 59 as an upper limit (Fig. S4).

In the following analysis, all classification experiments were performed with the 59-mer neighborhood context. (For detailed AUC statistics please refer to Tables, S5, S6, S7 and S8.)

### Incorporating GC% feature did not improve the classification performance

As described in the introduction, the existence of a correlation between sequence GC% and mutation processes in mammals has been known for some time. We therefore considered whether inclusion of GC% as a feature would improve classifier performance. GC% was estimated from ±500bp flanking each mutation. Only the naïve classifier (M) performance was improved by inclusion of the GC% feature (Fig S5). The impact on classifiers containing sequence features ranged from no effect (M+I) to substantially worse (FS). We speculate that the improvement of M+GC% over the M feature set arises because the GC% term indirectly measures the base composition of the immediate neighborhood captured by the I term.

### Applying classifiers to the whole genome

From the classifier development process described above, we selected the LR classifier with *k* = 59, M+I+2Dp feature set, and hyperparameters *ℓ*_1_, *C* = 0.1 trained on the ~ 16, 000 data sample from chromosome 1. We applied this classifier to all mouse point mutations and display the results by chromosome in Fig S5. The vertical axis is the AUC score for all chromosomes. We distinguish chromosome 1 because it was used for training (see Materials and Methods). Typically, classifier performance on data on which it was trained is expected to be greater. From chromosomes 2-19 X, and Y, the mean and standard deviation of the chromosome AUC scores was 0.84 and 0.01 respectively. Thus, the LR M+I+2Dp classifier has a relatively good performance across the entire genome. Despite it’s markedly superior learning curve, the XGBoost genome classification performed only marginally better than the LR classifier (Fig S6).

### Performance of the one-class classifier was substantially worse

We sought to evaluate whether the mutation motifs associated with spontaneous mutations were sufficiently distinctive as to allow a machine learning algorithm to effectively identify non-spontaneous mutations. This corresponds to an outlier analysis. We tackled this using a one-class (OC) Support Vector Machine (SVM). (See Materials and Methods for more detail.) We considered the same feature set choices as for the LR models in a 7-mer context. As shown in Fig S6, the M+I+2D feature set showed the best performance. However, all OC classifiers had much lower 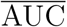 than even the simplest two-class classifier (M). Furthermore, the OC M+I+2D classifier applied to the entire genome exhibited a systematically lower AUC compared to the LR classifier (Fig S7).

## Discussion

We have sought to establish the extent to which the etiological relationship between flanking sequence and mutagenesis can be used to identify the mechanism via which individual point mutations originate. Genetic variants in the mouse arising from application of ENU, a potent chemical mutagen, were contrasted with those arising spontaneously. We show that ENU-induced point mutations are very strongly associated with neighboring bases in a manner that differs to their spontaneous counterparts. A two-class classifier performed markedly better to the current standard technique for identifying ENU mutations and was robust to genomic sequence attributes that have previously been shown to affect mutation processes. Our examination of the potential for machine learning based on the single category of spontaneous germline mutations revealed substantial challenges remain to resolving this more general case.

Comparison of the mutation spectra between spontaneous and ENU-induced germline mutations supported previous conclusions. The spectral analysis compared the breakdown of mutations from a single base mutation from ENU-induced and spontaneous mutations. The proportions of A→G* and A→T* mutations were substantially increased ~1.5 fold and ~7.5 fold, respectively, in the ENU-induced compared to spontaneous sample. These observations are consistent with previous reports (Barbaric et al., 2007; Takahasi et al., 2007; Justice et al., 1999; Noveroske et al., 2000). The abundance of A→G* point mutations in *both* the ENU-induced and spontaneous samples underscores the challenge of using mutation direction alone for classifying mechanistic origin, and the likelihood that such an approach will be error prone.

Our analyses established that the DNA sequence flanking ENU-induced mutations does contain distinctive information. After correcting for multiple hypothesis tests (Holm, 1979), highly significant associations between neighboring bases and point mutations were found for the ENU-induced sample, along with highly significant differences in neighborhood between the ENU-induced and spontaneous mutations. As ENU induces an elevated rate of DNA lesion formation, it seems plausible these differing neighboring base associations reflect that chemistry. Alternately, they may derive from operation of different DNA repair processes to those typically active in the germline (Noveroske et al., 2000; Takahasi et al., 2007; Shrivastav et al., 2010). In addition to the independent neighborhood effects, all ENU-induced mutations were found to be significantly associated with higher-order effects. Similar to what we was observed from humans (Zhu et al., 2017), the higher-order effects on ENU-induced mutations were evident in a manner such that bases at physically contiguous positions showed the largest RE (Fig S2). The latter may reflect the importance of base stacking on helix stability (Yakovchuk et al., 2006)

The analyses of the influence of sequence neighborhood on ENU-induced point mutations clarify previous reports. (Barbaric et al., 2007) found a significant enrichment of base G or base C at one of the two most immediate flanking positions. Their measurement encompassed all 12 mutation types and thus could not resolve whether this was a systemic influence of ENU, or one related to a specific point mutation. Indicating it is the latter, the results identified this specific pair of neighboring bases as highly significantly associated with ENU-induced A→G*. This result contradicts the claim, by (Bauer et al., 2015), that there are no neighboring base influences. We note that those authors did not formally test this hypothesis.

A succinct LR model was capable of strong performance, even when trained on just a small fraction of the total data. The current standard classifier, model M, represents the baseline performance. M considers only mutation direction and ignores sequence neighborhood entirely. The performance 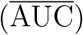 of the M+I+2D feature set on the trading data from mouse chromosome 1 was ~ 7% better than that of M while the fully saturated (FS) model exhibited comparable performance (Fig S2). This observation indicates that including dependent effects with order > 2 confers little benefit to classification performance. This observation is consistent with the results from our log-linear analysis which showed a small residual deviance after fitting the I+2D model (Fig S2).

The GC% statistic, previously correlated with mutation processes in mammals, was determined to be a crude surrogate of more explicit neighborhood features. GC% is a sequence composition summary statistic. Inclusion of this feature in the classifier only improved the M model. In all other cases it had no effect or reduced classifier performance (Fig S5). This result emphasizes the mechanistic role of individual bases, as reflected by the mutation motifs (Fig S1), rather than a more general property (e.g. the local DNA melting point) of a sequence region.

Application of the developed LR classifier to the whole genome produced a greater performance than what we observed on the training chromosome. We evaluated classifier performance on a per-chromosome basis to facilitate evaluating whether a relationship existed between classifier performance and the distinctive *k*-mer distributions reported for mammal sex chromosomes (Huttley et al., 2000). For the LR classifier, the AUC from the combined sex-chromosomes was the lowest of all AUC scores. However, it was not markedly distant from the range of autosomal values, indicating the discriminatory resolution of the LR classifier was largely robust to such differences.

It is worth noting that our LR classifier was trained using relatively balanced data, that is the number of ENU and germline mutations were comparable in our data set. This design reflects our interest in understanding what sequence factors affect classifier performance, rather than the specific objective of delivering a classifier for studies employing ENU. In such studies, the mutation classes will be highly imbalanced as we expect many more ENU than spontaneous mutations (up to 100-fold excess). This attribute needs a different trade-off between false positive and false negative predictions from the classifier. There are several extensions to this work that may be useful when a practitioner attempts the class imbalanced task. First is to consider using a performance metric that is less sensitive to class imbalance (Davis and Goadrich, 2006). Second is to extend the learning method to manage class imbalance during both the training and prediction steps. This can be done in part using re-sampling methods or cost sensitive methods (e.g. Haixiang et al., 2017). Third is to consider the suitability of the classifier for the imbalanced case. The categories of classifiers differ in the ease with which they can represent class imbalance. NB classifiers explicitly accommodate such imbalance via class priors. Application of LR for classification on imbalanced data is less obvious, although approaches involving adjustment of the intercept terms have been proposed (King and Zeng, 2001). The different performance of these two classifier categories evident in this study (discussed further below), however, indicates a final choice for the imbalanced case requires further investigation.

A one-class classifier would also provide a means for generic identification of mutations that did not match a designated reference sample. For instance, a forward genetics screen employing ENU where spontaneous mutations are rare. While the outcome of feature selection identified the feature set M+I+2D as the best performing OC classifier, the 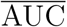 from the genome was 0.67. This is significantly better than a random guess, but just ~84% of the two-class classifier performance. This discrepancy in performance likely reflects the overlap between sequence features of the ENU-induced and spontaneous mouse germline mutations. Since the one-class models are trained only on one sample, they are very sensitive to irrelevant neighborhoods compared to the two-class classifiers. In other words, the presence of “noise” makes it difficult to identify neighborhoods that are unique to the positive class. Furthermore, the one-hot encoding (see Materials and Methods) for one-class classification produces a sparse table for the sample size, which can reduce classification performance.

Both the choice of *k* and the corresponding feature set had a pronounced impact on the results obtained here. For values of *k* in {3, 5, 7}, we considered the full set of alternate feature sets, i.e. M, I, and all possible dependent interaction terms. Classifier performance increased with value of *k* but a trade-off between classifier performance and memory usage precluded naïve extension to large *k*. Consideration of the simpler M+I model for much larger values indicated potentially quite substantial gains in performance may be attainable. Learning curve analysis of the M+I model for *k* = 59 returned 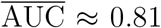. Further extension of this model was restricted to 2D terms between proximal positions. This M+I+2Dp model further increased classifier performance to 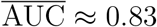 (Fig S4).

The generally poorer performance of the NB approach (Fig S4) led us to discard it. There have been systematic examinations of differences between LR and NB classifiers (Ng and Jordan, 2002). These differences are due to the different structural assumptions used by the classifiers. LR is a discriminative classifier, and it directly estimates the conditional probability of interest. NB is a generative classifier, estimating both the prior and likelihood before using them to estimate the posterior probability of interest. The design choice of estimating the likelihood makes NB more sensitive to data that violates the Gaussian noise assumption. Therefore when the underlying data does not exhibit Gaussian noise, LR classifiers have lower asymptotic error than NB. In addition, if training sizes are relatively large, then LR performs better than the NB classifiers (Ng and Jordan, 2002).

Our results have established the utility of including a representation of sequence neighborhoods in classifiers for resolving point mutation origins. There remain open questions as to why should large *k* be so informative, when the analysis of information content of neighboring bases revealed a quite restrictive limit (Fig S3 and Zhu et al., 2017). Perhaps, as speculated previously (Bauer et al., 2015), this reflects broader sequence features correlated with open chromatin status during spermatogenesis. Irrespective of biological mechanism, the marked improvement in classifier performance we were able to achieve is suggestive further improvements are possible.

We have shown that neighboring positions can be used to classify the mechanistic origins of mutations using machine learning techniques. The LR classifier can be expressed in relation to the log-linear models and this relationship allowed us to “dissect” the contribution level between different positions. However, the classifier features used here were mainly designed for two classes. While we used them for the one-class classification as well, and the performance was better than random guessing, the best customization of feature selection for the one-class classifier remains unresolved.

## Materials and Methods

### Spontaneous and ENU-induced germline mutation data

We constructed the data set for mutation origin identification from Ensembl release 88 and an ENU variation database from the Australian Phenomics Facility. The number of variants per chromosome are reported in Table S4 in the Supplementary Information.

As defined in the Introduction and Results sections, we adopt the following notation to refer to the 12 different point mutations. The mutation of base X into base Y is indicated by X→Y. We denote a point mutation and its strand complement using *. For instance, A→G* refers to both A→G and its strand complement T→C.

### Mouse spontaneous germline variants

The germline spontaneous variant data was obtained from the Ensembl database using EnsemblDb3 (http://ensembldb3.readthedocs.io). For each genetic variant we obtained the SNP name, genomic location, effect and alleles. Only biallelic SNPs were used. Because the Ensembl database did not include mutation direction for mouse variants, we computed mutation direction using phylogenetic methods.

Inference of mutation direction was performed using ancestral sequence reconstruction (Yang et al., 1995). The genomic alignments of mouse protein coding genes and their one-to-one orthologs from the rat and squirrel were sampled from Ensembl using EnsemblDb3. Checks were performed to ensure the obtained syntenic alignments could be used. Specifically, only mouse genetic variants where the genomic alignment contained unambiguous bases for all species were retained. The genomic alignments were sliced to be centered on a genetic variant. We fitted the HKY85 substitution model (Hasegawa et al., 1985) by maximum likelihood using PyCogent3 (Knight et al., 2007, http://cogent3.readthedocs.io) and estimated the most likely base at the mouse variant locus for the common ancestor of mouse and rat. This ancestral base, which matched one of the reported mouse alleles, is taken as the starting base and this allows inference of the mutation direction that produced the genetic variant.

A total 254,680 validated mouse germline spontaneous variants within protein coding regions were sampled. These variant records are further separated into sub-categories according to mutation direction and chromosomal location (Table S4).

### ENU variants

ENU induced variant data examined in this study were obtained from the Australian Phenomics Facility website (https://pb.apf.edu.au/phenbank/download/). In the database, each genetic variant record includes the variant identifier, genomic location, putative effect, reference base and variant base. The mutation direction is inferred as a change from the reference to variant base. Only synonymous and non-synonymous mutations in mouse exonic protein coding regions were used for this study. This resulted in 234,177 ENU-induced mutations. Summary details of ENU variant records regarding mutation direction and the chromosomal location are presented in Table S4.

### Association of neighboring bases using log-linear modeling

We employ our previously published log-linear methods (Zhu et al., 2017) and corresponding MutationMotif software (https://github.com/HuttleyLab/MutationMotif) for evaluating the association of neighboring nucleotides and spontaneous and ENU-induced point mutations in the mouse. In summary, these methods allow statistical evaluation of the association between point mutations and bases at individual, or multiple, sequence positions. They further allow comparisons between samples for these associations. The log-linear models operate via comparing the count of observed bases at a position in sequences for which the point mutation is known against a paired reference distribution of counts from unmutated sequences. The association of bases at a single position with point mutations is referred to as an independent effect and the influence of bases at two or more positions are referred to as dependent effects. These tests were used to assess the null hypotheses that ENU-induced point mutations occur independent of neighboring bases. We also tested the null that the neighboring base effects were the same between ENU-induced and spontaneous point mutations.

Mutation motifs were visualized in a sequence logo style. The stack height in these figures corresponds to relative entropy (RE). Individual letter heights within a stack represent the relative magnitude of the residual from the log-linear model for that letter. Base(s) that are overabundant in mutated sequences are on top with a normal orientation. Base(s) with letters rotated 180° are underrepresented in mutated sequences.

### Prediction of mutation origins

A difference in the association of neighboring bases with spontaneous and ENU-induced mouse point mutations provides a basis for using machine learning classifiers to predict mutation origin. We consider two scenarios for such analyses. In the first, two mutation classes are known in advance allowing development of a discriminating function. In the second, we consider the case in which only one mutation class is known in advance and we seek to identify mutations that are ‘outliers’ to this known class. Of the numerous alternate machine learning techniques that could be applied to the two-class problem, we employ logistic regression (hereafter LR), XGBoost (hereafter XGB, and Naïve Bayes (hereafter NB). We employ LR because of its similarity to the log-linear modeling approach described above. XGBoost was chosen as a representative of ensemble style learning algorithms. NB was chosen as it is methodologically quite different from LR and has also been used extensively for sequence classification. For the one-class problem, we use a support vector machine (SVM). We use the open source software library scikit-learn (Pedregosa et al., 2011) for these along with the XGBoost library (Chen and Guestrin, 2016).

### Logistic Regression

The parametric nature of LR facilitates mechanistic interpretation of the developed classifier (Wålinder, 2014; Prosperi et al., 2009). This is of particular interest here as we seek to relate attributes of the biological data to classifier performance. LR is based on the logistic function (James et al., 2013) as shown in Eq 1. The response value of LR ranges from 0 to 1. In classification, the probability that an observation belongs to a certain mutation class (e.g. ENU) is expressed in Eq 2. We classify mutation *X* as originating by mutation class 1 if *Pr*(*Y* = 1|*X*) is greater or equal to 0.5.

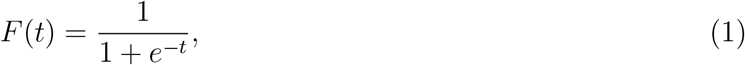

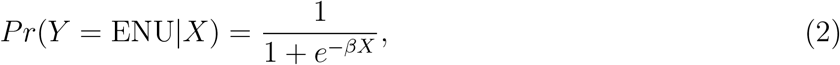

The approximate probability *π_q_* of a mutation given feature sets can be expressed as:

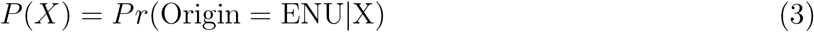

*P*(*X*) ranges between 0 and 1, and the logistic regression expression of *P*(*X*) is

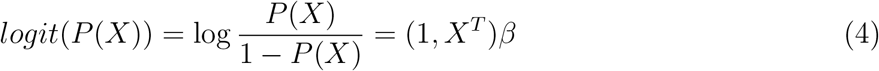

or

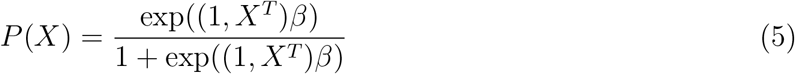

where *X* is the input vector of features. *β* is a parameter weight vector describing how important each feature is, a larger *β* value indicating a more important feature, however, a large *β* may also indicate that the associated feature is over-fitted. Also, according to Eq 5, we found that different settings of *β* value will lead to different prediction probability. We want our classifier to perform as accurate as possible, therefore, we need to find the optimal set of *β* which generates the maximum prediction probability without over-fitting feature weights. The *ℓ*1 norm (*ℓ*1) regularization was performed to achieve this.

In this study, we used *ℓ*1 regularization because it prunes out unneeded features by setting their associated weights to 0. This characteristic allows us to understand the contribution of each feature better. Mathematically, *ℓ*1 regularized logistic regression by solving the following optimization problem (Pedregosa et al., 2011)

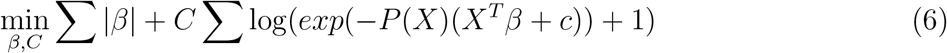

where hyperparameter *C* is a positive constant that balance how much we care about fitting the training data compared to penalizing large weights. *C* was tuned during cross validation process to maximize the likelihood, and the according estimates of *β* were stored for subsequent use in predicting mutation origin based on the selected feature set.

### Naïve Bayes

NB classifiers are built upon the assumption of conditional independence of the predictive variables given the class. This assumption is typically violated. However, as our variant data were randomly sampled from different mice the dependency between mutations is relatively low and thus the NB classifier was expected to perform reasonably.

To learn information from training samples according defined feature sets and to predict origins of mutation with NB classifier, similar to the logistic regression classification, each variant data is ultimately represented as a vector of binary features including mutation direction and the neighborhood sequences. In a NB algorithm, the posterior probability a variable was ENU-induced given a feature set is calculated as

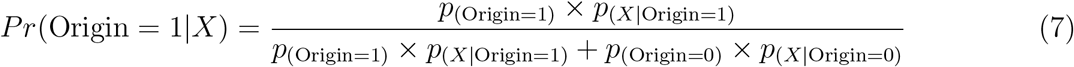

where Origin classes 1, 0 correspond to ENU-induced and spontaneous germline mutations respectively. This product goes over all data in the training sample, where *x_q_* represent feature vectors. If the resulting posterior probability is higher than a defined cutoff threshold, then a mutation is classified as ENU-induced mutation; otherwise, it is considered to be a normal mouse germline mutation. To optimize *Pr*(Origin = 1|*X*), key components *p*_(*X*|Origin)_ for each origin class, in Eq 7 is estimated by a smoothed version of maximum likelihood

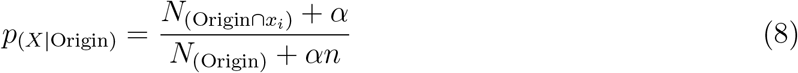

where, for each origin class, 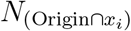 is the frequency count of feature *x*_*i*_, *x*_*i*_ ∈ *X* appearing in a sample belonging to that particular origin class, and similarly, *N*_(Origin)_ is the frequency count of sample belonging to a particular origin class. *α* is the smoothing factor and value of *α* is tuned during cross validation process to optimize the result, and *n* is the number of features.

One of the main advantages of NB classifiers is that they are probabilistic models. In addition to predicting the class label of a point mutation can be predict, the probability of the class labels is also generated.

### Gradient boosting using XGBoost

Gradient boosting is an ensemble class of machine learning algorithms with XGBoost a popular variant (Chen and Guestrin, 2016). Ensemble learning approaches evaluate and combine multiple base functions for classification. In XGBoost, the base functions are classification and regression trees, which are additively combined using boosting. The objective function used in boosting uses logistic loss (the same as logistic regression) and a penalty term involving the complexity of the trees. Gradient boosting techniques operate such that the function that most improves the overall score is added at each iteration.

We address the challenge of training XGBoost by an incremental search over parameter space. We specifically employed https://github.com/Jie-Yuan/xgboost-tuner (version 0.1.2) to train XGBoost classifiers. This implements a best practice approach to exploring the numerous possible settings that tune how the classifier training is done the parameters that affect performance. We specified logistic regression as the objective function and employed the incremental (exhaustive) grid-search with 3-fold cross validation. The exact scope of the parameter grid used for the incremental search is specified within the `mutation_origin.classify.xgb` function.

### One-class classification using SVM

The logistic regression classifier and Naïve Bayes classifiers are designed to solve the two-class situation, that is to distinguish whether a mutation is a germline spontaneous mutation or an ENU-induced mutation. An interesting possibility is that may arise in real studies is that the properties of an alternative mutation mechanism are unknown, but a well characterized reference data set exists. In the case, we are interested in finding out whether a mutation is likely to be a member of the reference set. In the present case, the reference distribution corresponds to spontaneous germline point mutations and we wish to know whether we can successfully identify the ENU-induced mutations.

To address this question we employed a one-class SVM algorithm to identify whether or not a mutation is considered to be a spontaneous mutation given training data and a proposed feature set. The spontaneous mutations are now the target objects and are labeled as +1, and the ENU-induced mutations are outliers and are labeled as −1. Training of the one-class classifier involves analysis of only spontaneous mutations to learn a classification boundary. To make the one-class SVM classifier results comparable to the logistic regression classifier results, we adopted the linear kernel when constructing the classifier, and we have the following decision function

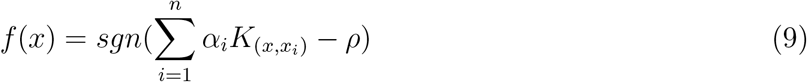

where *α*_*i*_ are the Lagrange multipliers, *ρ* are the parameters of the hyperplane, and 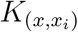 is the linear kernel function. The classifier are then applied to the test data to determine whether a mutation is a spontaneous mutation (positive) or a ENU-induced one (negative).

### The feature sets employed for classification

The machine learning approaches require numerical representation of the data. The choices of features employed will affect the final performance of a classifier. If the feature is not enough to describe a data sample, then there is not enough information available for a classifier to learn the data structure well. Intuitively, increasing the number of non-correlated features typically increases classification performance. However, if too many features are selected, it is computationally expensive.

We explored four different types of features: mutation direction, independent neighborhood effects, dependent neighborhood effects, and GC%. Mutation direction, which we represent by M, is the point mutation direction (e.g. C→T), of which there are 12 possible. Independent effects, which we represent by I, is the influence of bases at flanking positions independent of what bases are present at other positions. Dependent effects are indicated by #D where # is the effect order. For example, a second-order dependent effect, represented by 2D, is the influence of the bases at 2 separate positions. For a 5-mer with the mutation at the central base there are 6 possible pairs of positions. The fully saturated feature set, represented by FS, contains the mutation direction and all possible independent, dependent features. We further employ a restriction on the dependent effect, that the component positions were proximal to each other in the sequence (after excluding the mutated position). We represented this feature set variant using a “p” suffix, e.g. 2Dp. For a 5-mer, there are 3 2Dp features. Each of these features are logical propositions which are represented by a one-hot encoding (illustrated in Table S2).

We further considered the percentage of G and C nucleotides (GC%) around a point mutation. We include this property as a significant positive correlation exists between inferred mutation rate and GC% in mammals (Hodgkinson and Eyre-Walker, 2011). The GC% is obtained from 500bp flanking sequences around a mutation (500bp from each side), numerical data.

For feature sets that were strictly categorical, genetic variant data was encoded with the one-hot encoding scheme. We use a {+1, −1} encoding for binary features, where +1 indicates that the logical proposition is true and −1 indicates that the logical proposition is false. Application of this process is illustrated for a small example in Table S2. In this example, the first record was derived from ENU-mutagenised mice and for the feature Variant class, is assigned +1 for the ENU value, −1 for the Spontaneous value. This process continues such that for a single record, only one of the possible values of a feature can be assigned +1.

As the GC% feature is not categorical, a different numerical representation was employed. The mutation direction features are categorical features, and labeled as +1 if true, or −1 if not true. On the other hand, the GC% feature is a numerical feature requiring a numerical representation of average GC percentage in neighboring sequences around a mutation, ranges from 0% to 100%. Because the range of values of raw data varies widely, the proposed classifier may not work properly without normalization. During a normalization application, the different numerical scales of GC% and the one-hot encoded categorical feature values were adjusted to a notionally common scale. This leads to these different features having approximately the same effect in the computation of similarity (Aksoy and Haralick, 2001). We used the scikit-learn StandardScaler to obtain a scalar for a normalized transform of the training data. The scalar derived from the training set was also used to normalize the test data.

### Machine learning experimental design

There are multiple factors that may influence the performance of a classifier. These include the choices regarding the algorithm, the values of associated hyperparameters and the feature set to be used for classifying. In addition, there are design considerations concerning selection of data for training and subsequent testing. The processes we employed for both the one- and two-class classification problems are illustrated for LR, NB and OC in Fig S8. Our core algorithm choices are described above. Our experimental design involved training our classifiers on data derived from mouse chromosome 1 only. For each algorithm, we used cross validation to tune the hyperparameters and optimize the classifier. For every cross validation iteration, we firstly perform a random train-test split, and divide our data sets into training data and testing data. Then inside the training data, we further split training data to actual training data and validation data (Fig *S*9). We train the classifier on training data, set hyperparameters on validation data and finally evaluated classification performance on testing data. Within each validation process, we compared algorithm performance with different hyperparameter values and the hyperparameter generating the best performance for the available data was saved. For each classification experiment, this process was repeated 10 times.

For the LR classification, the hyperparameter *C* is the trade-off regularization parameter which trades off misclassification of training examples against simplicity of the decision surface. A low *C* makes the decision surface smooth, while a high *C* aims at classifying all training examples correctly by giving the model freedom to select more samples as support vectors. We considered candidate *C* options from the log-scale of: 0.01, 0.1, 1, 10, 100. The *C* value that resulted in the best performance (please refer to Classifier performance evaluation section), was chosen for all subsequent analyses.

For the Naïve Bayes classification, the hyperparameter alpha is the Laplace parameter used to smooth categorical data. We considered candidate alpha options of: 0.01, 0.1, 1, 2, 3. The value of alpha which resulted in the best performance was chosen for all subsequent analyses.

### Classifier performance evaluation

We evaluated classifier performance using the area under the receiver operating characteristic curve (AUC). One of the advantages of using AUC score as the performance measure is that the score does not require choice of a cutoff threshold. Many binary classification algorithms compute a series of performance scores (e.g. overall accuracy, sensitivity, and specificity), and they classify based upon whether or not the score is above a certain threshold. Therefore, as the choice of threshold is of particular importance in these scoring schemes, shifting of the threshold may dramatically alter the score and thus the performance of a classifier. AUC score has the advantage of illustrating the trade-off between sensitivity and specificity for all possible thresholds rather than just the one that was chosen by the modeling technique. The AUC also has a probabilistic interpretation. Specifically, AUC is the probability that the predicted value (and thus rank) of a randomly drawn positive case is higher than the predicted value of a randomly drawn negative case. Here, the AUC scores of the different experiments are reported and we interpret a larger AUC score as indicating better classification performance.

### The effect of increasing the number of examples during training

The whole classification process is achieved by implementing training and testing phases. In the training phase, a set of data and their respective labels are used to build a classification model. In the test phase, the trained classifier is used to predict new cases. Overlap sampling between training and testing data will make the prediction performance of a classifier overly optimistic, because of the overfitting problem. To avoid the overfitting situation, for each experiment, to start with, both ENU-induced mutations and mouse germline mutations are split into two non-overlapping sets for training and testing.

The accuracy of a classifier improves with the number of observations used to train the algorithm. This improvement tends to be rapid initially, and then when the training size is sufficient to a point, the improvement decreases gradually. The “learning curve” is used to describe this phenomenon, and is used to estimate the number of samples needed to train a particular classifier to achieve its optimal accuracy (Mukherjee et al., 2003). To plot learning curves and find the desired training size, after selecting a specific classifier and set of features, we used progressively larger samples of observations to train the classifier and then plot accuracy performance against the number of training observations.

## Availability of data and materials

The pre-processed data used in this study are available at Zenodo https://zenodo.org/record/1204695 under the Creative Commons Attribution-Share Alike license. Data files are typically gzip compressed standard formats, e.g. tab delimited text files, fasta formatted sequence files. The source code for a command line application is made available under the BSD clause-3 license at https://github.com/HuttleyLab/mutationorigin and https://zenodo.org/record/3497585. The scripts used to perform the data sampling and analyses reported in this work along with the derived data are freely available at https://github.com/HuttleyLab/enuproject and https://zenodo.org/record/3497584.

## Acknowledgments

We thank B Kaehler, H Simon and H Ying for comments on earlier versions of the manuscript.

---

**Table S1:**
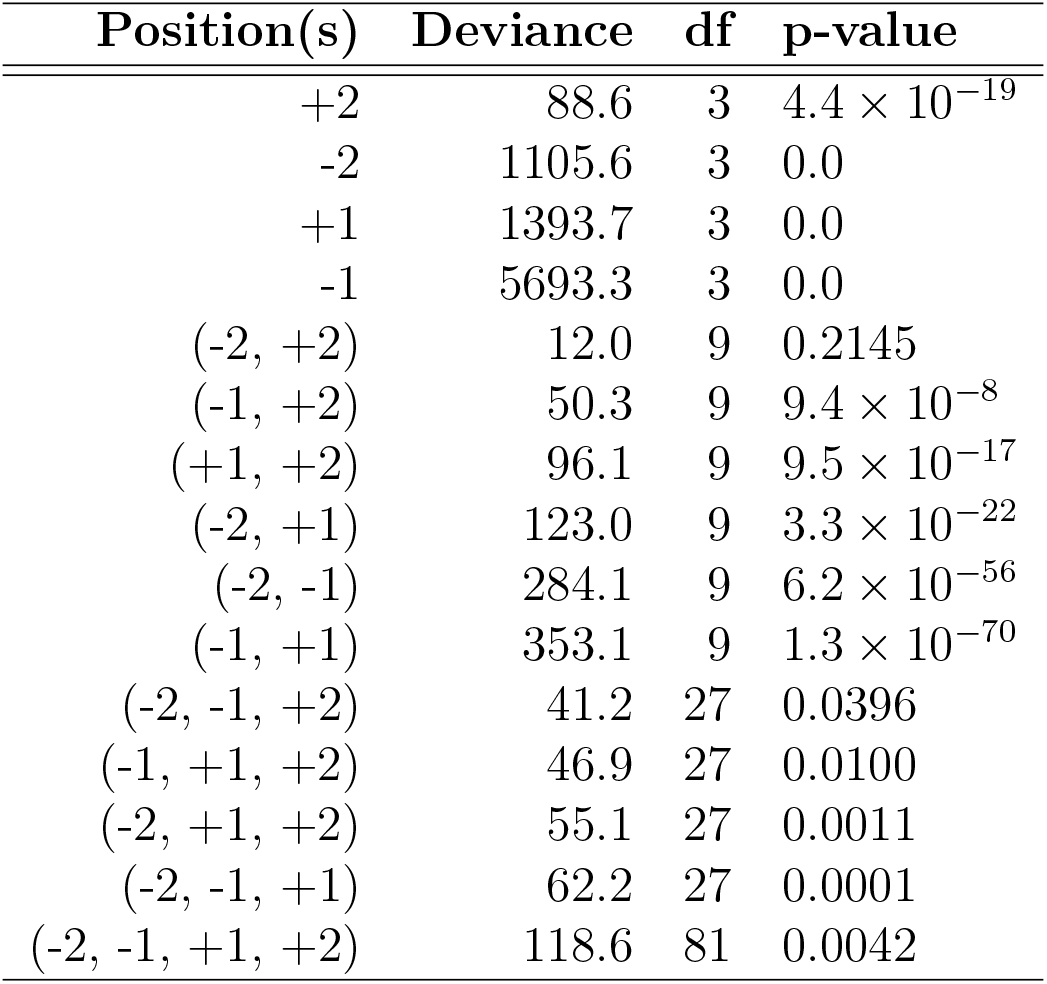
Log-linear analysis of mutation motif comparison between mouse germline and ENU-induced A→G mutations. Deviance is from the log-linear model, with df degrees-of-freedom and corresponding *p*-value obtained from the *χ*^2^ distribution.

| Position(s) | Deviance | df | p-value |
| --- | --- | --- | --- |
| +2 | 88.6 | 3 | $4.4 \times 10^{-19}$ |
| -2 | 1105.6 | 3 | 0.0 |
| +1 | 1393.7 | 3 | 0.0 |
| -1 | 5693.3 | 3 | 0.0 |
| (-2, +2) | 12.0 | 9 | 0.2145 |
| (-1, +2) | 50.3 | 9 | $9.4 \times 10^{-8}$ |
| (+1, +2) | 96.1 | 9 | $9.5 \times 10^{-17}$ |
| (-2, +1) | 123.0 | 9 | $3.3 \times 10^{-22}$ |
| (-2, -1) | 284.1 | 9 | $6.2 \times 10^{-56}$ |
| (-1, +1) | 353.1 | 9 | $1.3 \times 10^{-70}$ |
| (-2, -1, +2) | 41.2 | 27 | 0.0396 |
| (-1, +1, +2) | 46.9 | 27 | 0.0100 |
| (-2, +1, +2) | 55.1 | 27 | 0.0011 |
| (-2, -1, +1) | 62.2 | 27 | 0.0001 |
| (-2, -1, +1, +2) | 118.6 | 81 | 0.0042 |

**Table S2:**
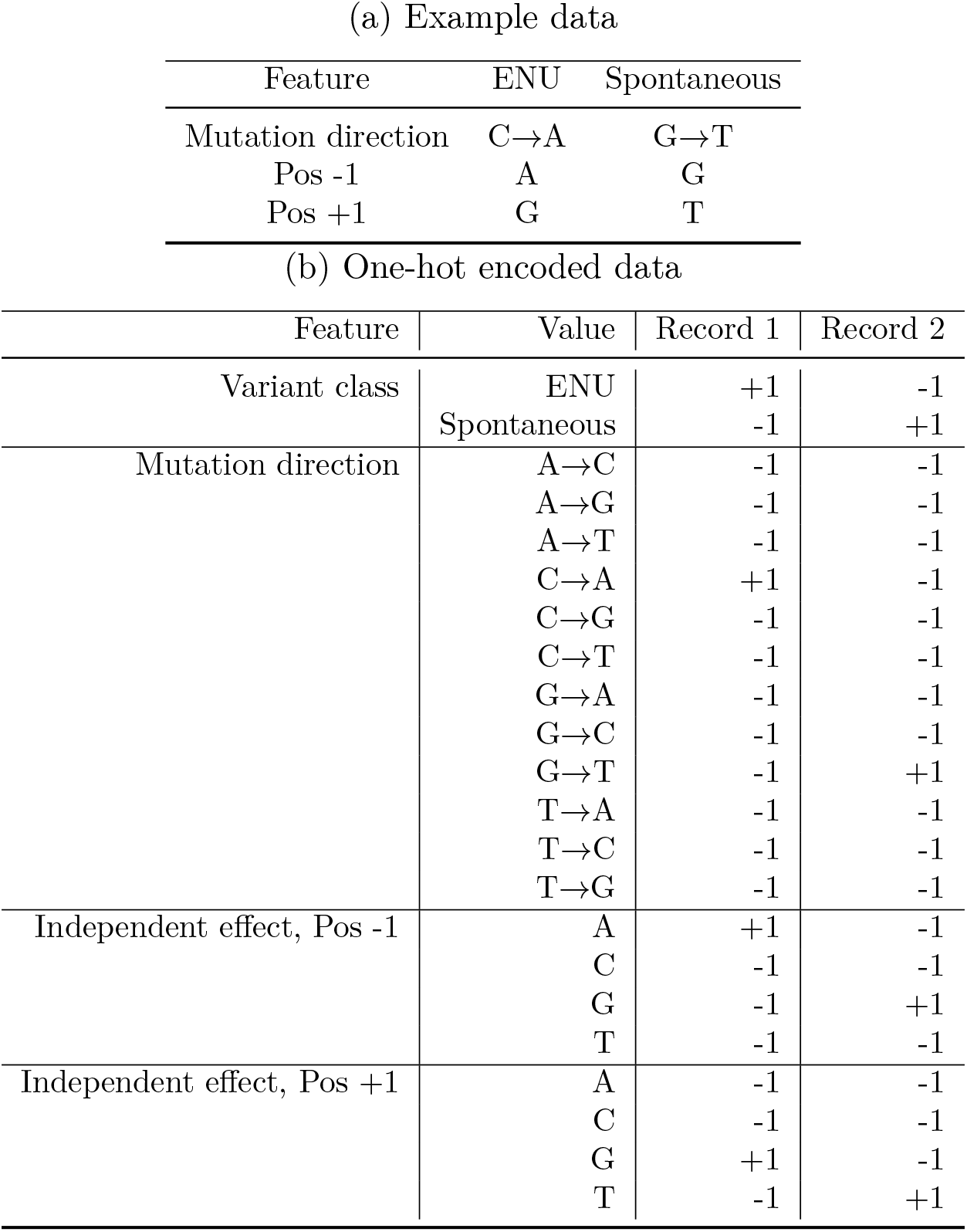
One-hot encoding of two mutation records for analysis. (a) An example raw data set containing an ENU and a Spontaneous mutation record. For each record, 1 bp neighboring bases on both side are shown (i.e. *k* = 3), positions −1, +1 are the left and right flanking neighboring positions respectively. (b) The one-hot encoding of the example data for a M+I classifier. In our notation, the feature ‘Mutation direction’ corresponds to M and the features ‘Pos’ correspond to I. Within a Feature, there are multiple possible values: 12 for the ‘Mutation direction’ feature, 4 for each ‘Pos’ features. For each record (column), only a single row within a feature can equal ‘+1’.

**Figure S1:**
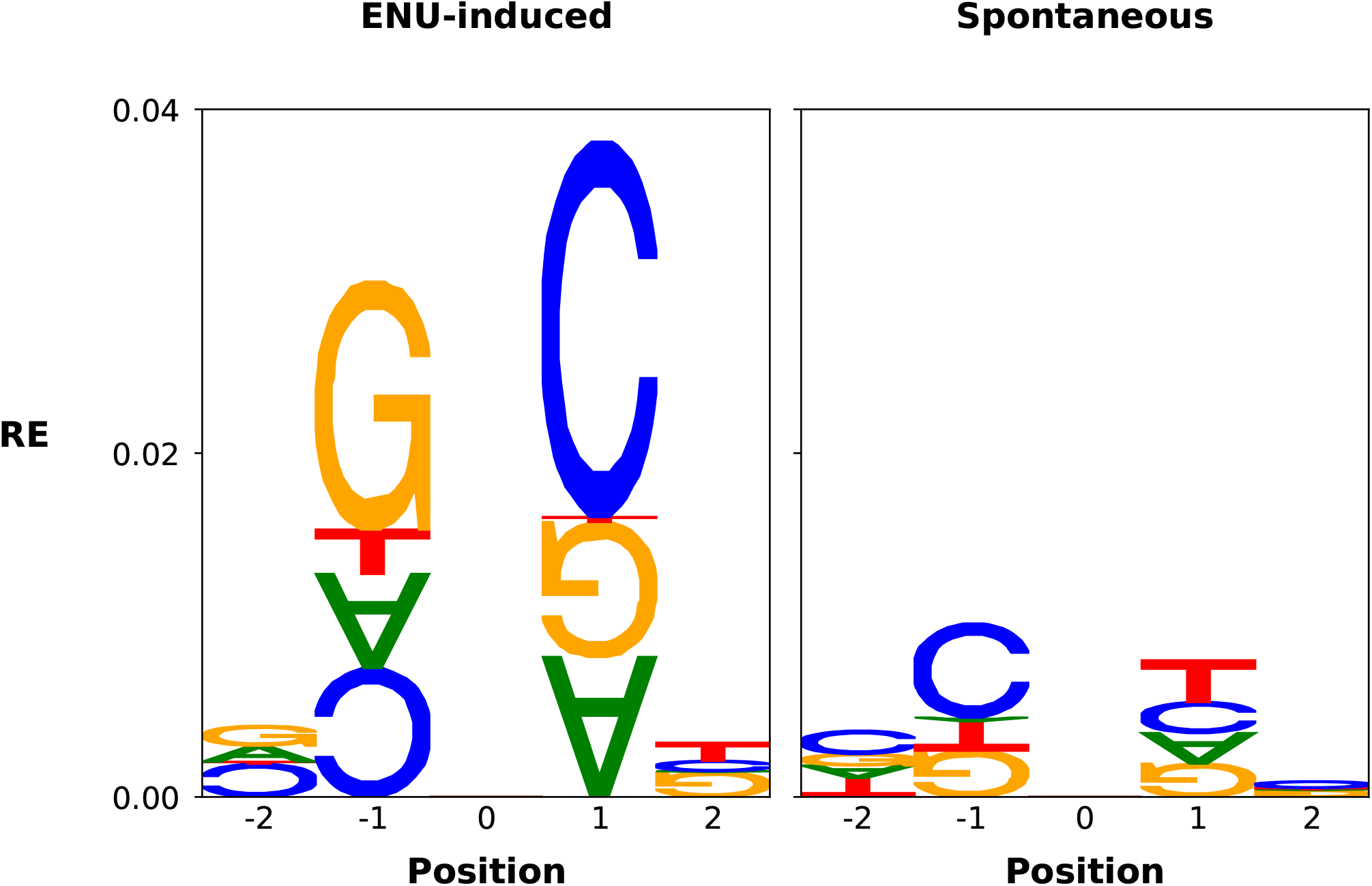
Neighboring base associations significantly differ between ENU-induced and spontaneous germline A→G mutations. Position is relative to the point mutation at position 0. RE is relative entropy, derived from the deviance of the log-linear model (Zhu et al., 2017). Letter height is proportional to the relative entropy term for that base. Normally oriented (180°-rotated) letters represent bases that are positively (negatively) associated with the point mutation. See Materials and Methods for more details.

**Figure S2:**
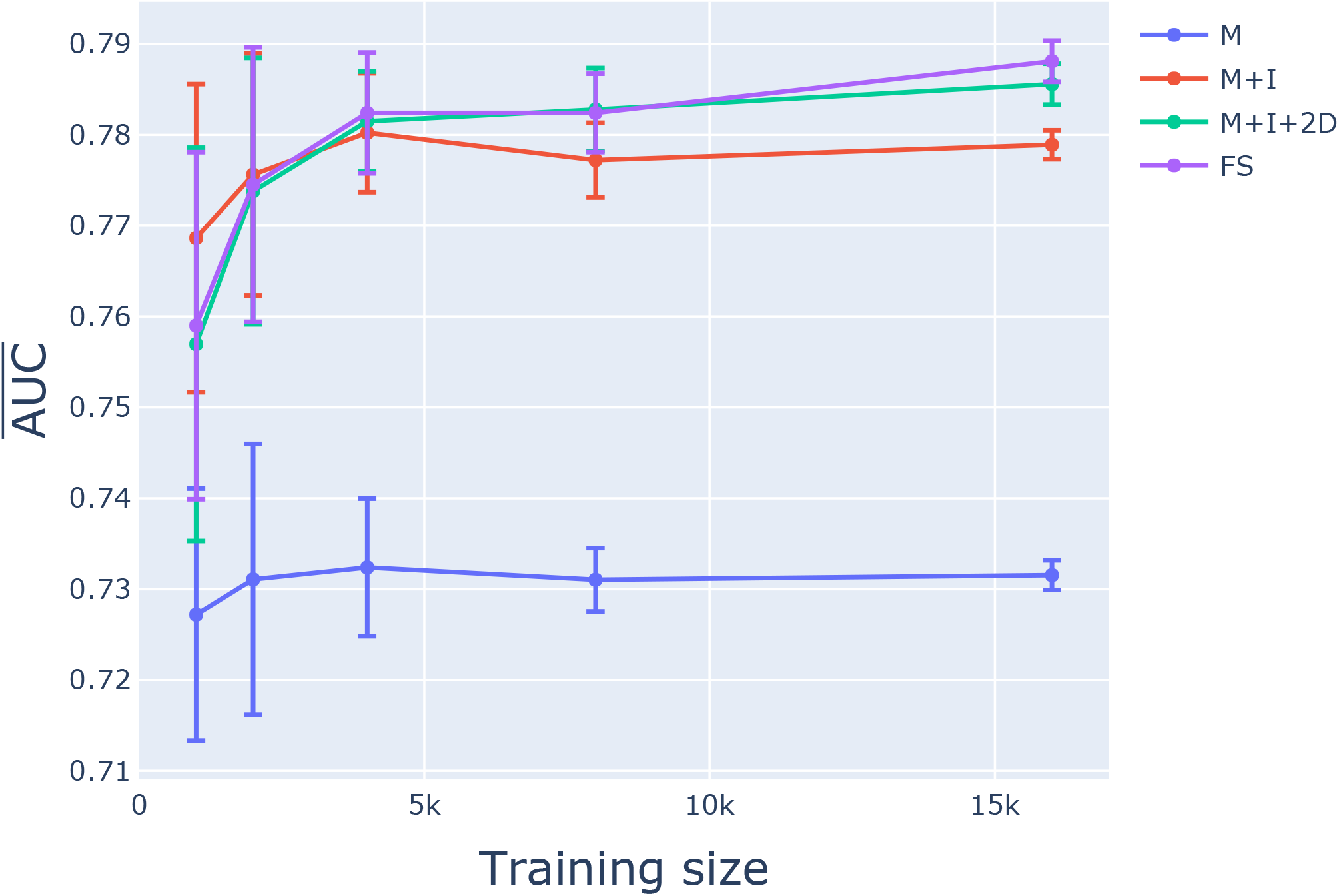
Model M+I+2D was sufficient for classifying mutations. Learning curves from training data are shown for four proposed classification models from 7-mers: M, M+I, M+I+2D and FS. The mean 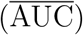 and standard error were calculated from the 10 chromosome 1 training samples. See the text for an explanation of model notation.

**Figure S3:**
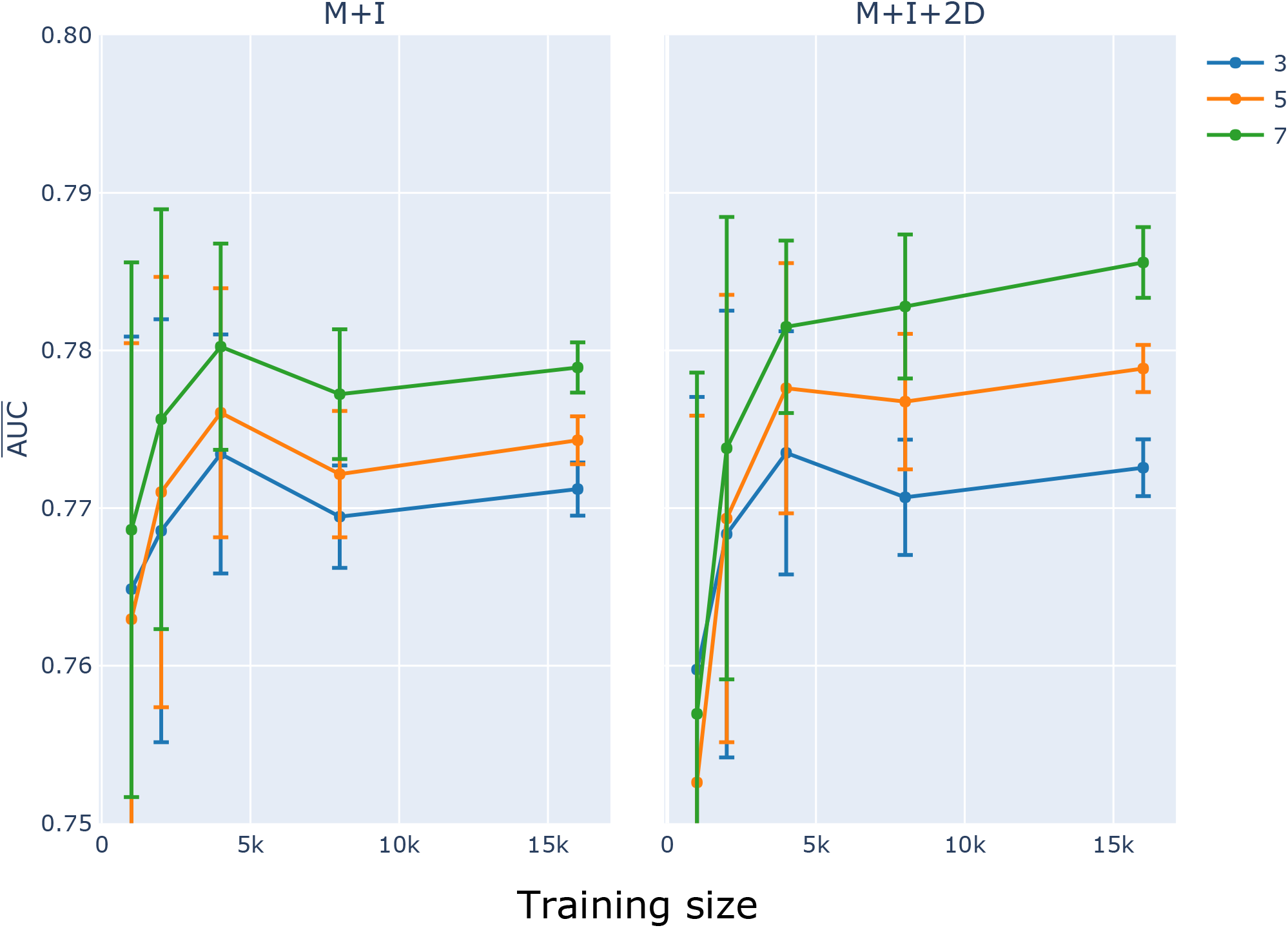
Classifier learning curves indicated increasing performance with *k*. The influence of *k*-mer choice on learning curves is shown for models M+I and M+I+2D. Plot titles indicate the model being evaluated. 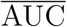 and the standard error were computed as described in Fig S2.

**Figure S4:**
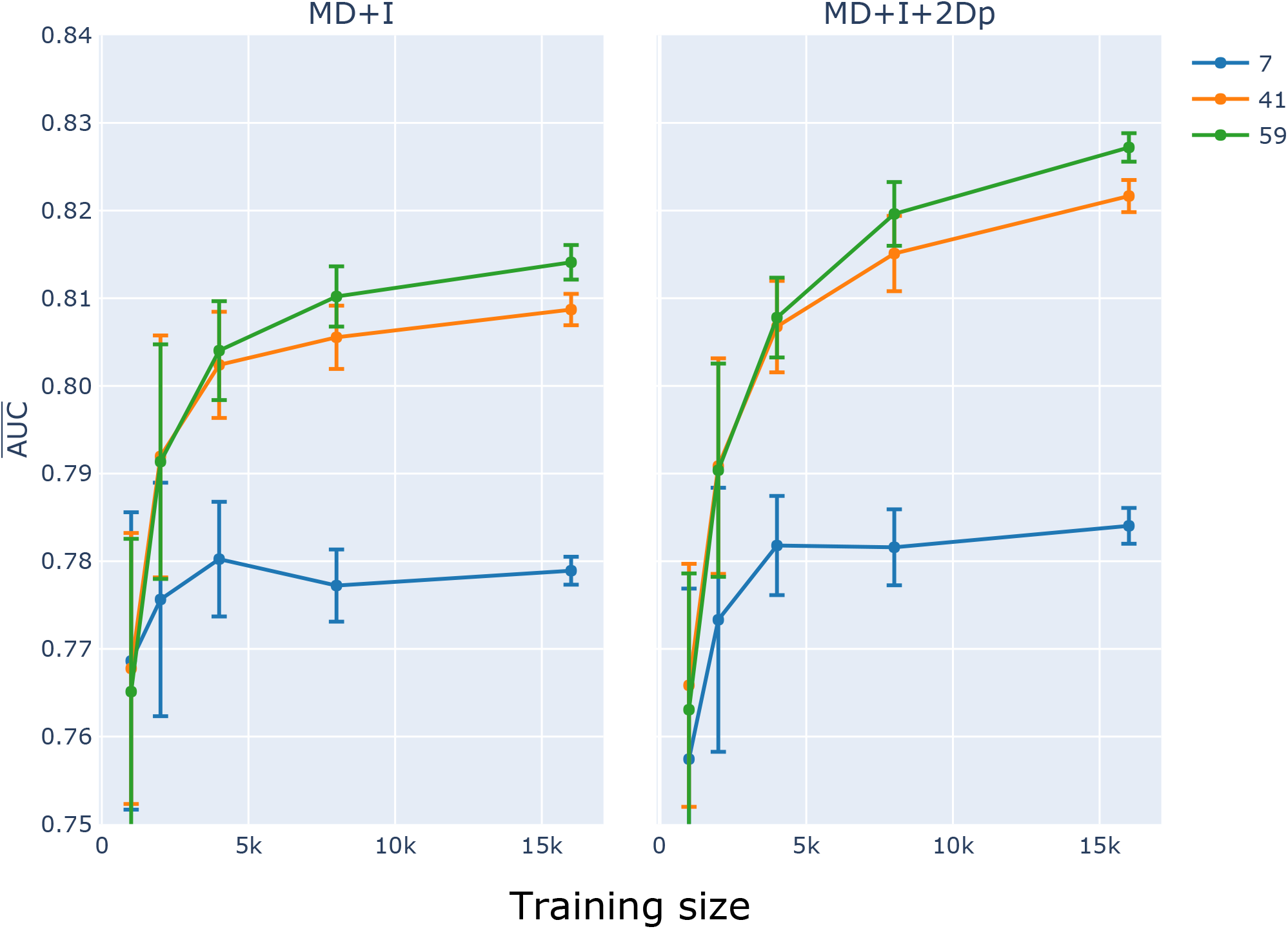
Large *k* and proximal 2D feature sets substantially improved classifier performance. Plot titles indicate the model being evaluated. 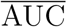 and the standard error were computed as described in Fig S2.

**Figure S5:**
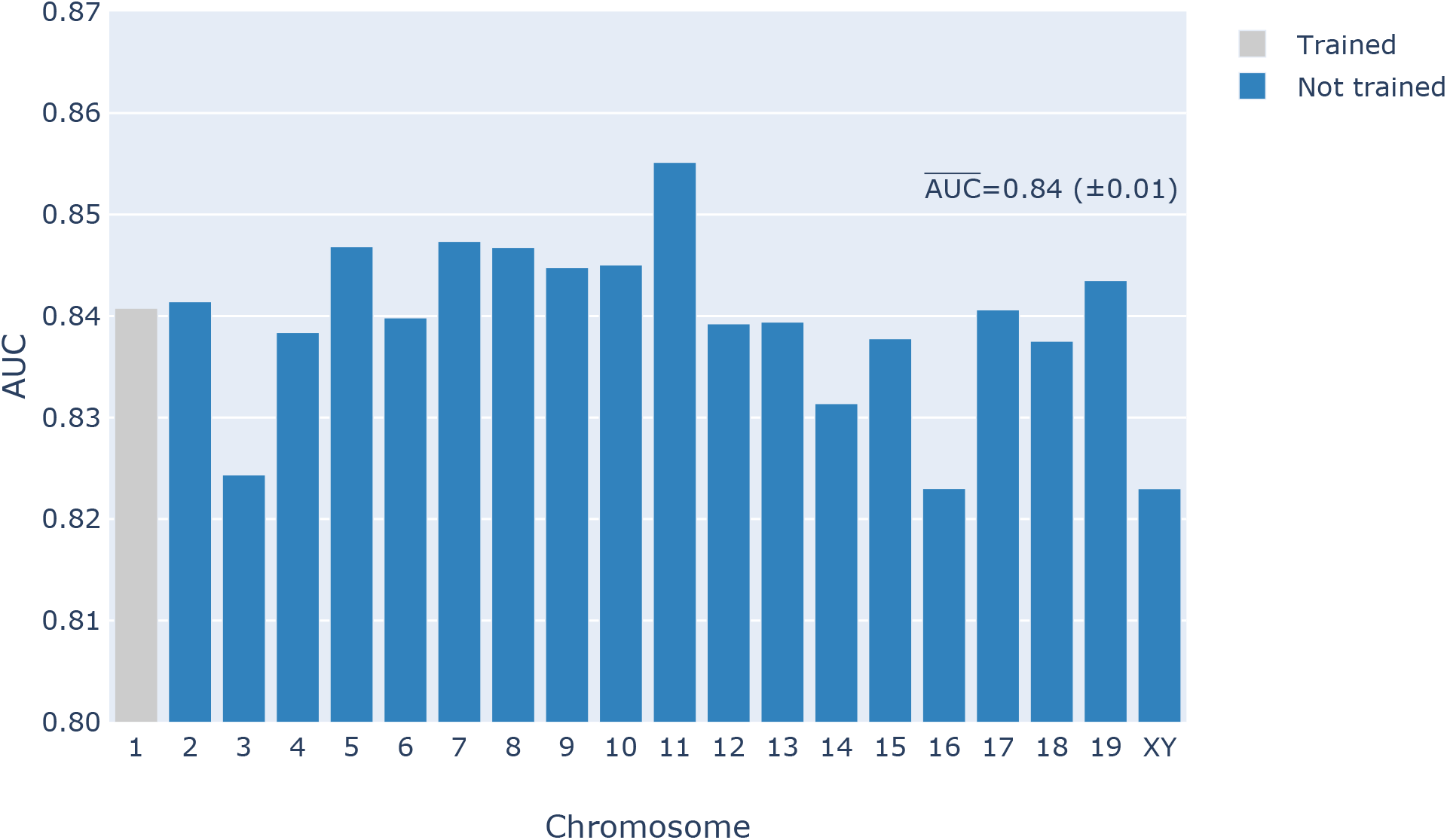
Per chromosome classification performance on the mouse genome of the best LR classifier. The classifier was trained on 16,000 mutations from chromosome 1 using a 59-mer M+I+2Dp feature set. The 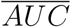 score from the chromosomes not used for training is shown on the figure.

**Figure S6:**
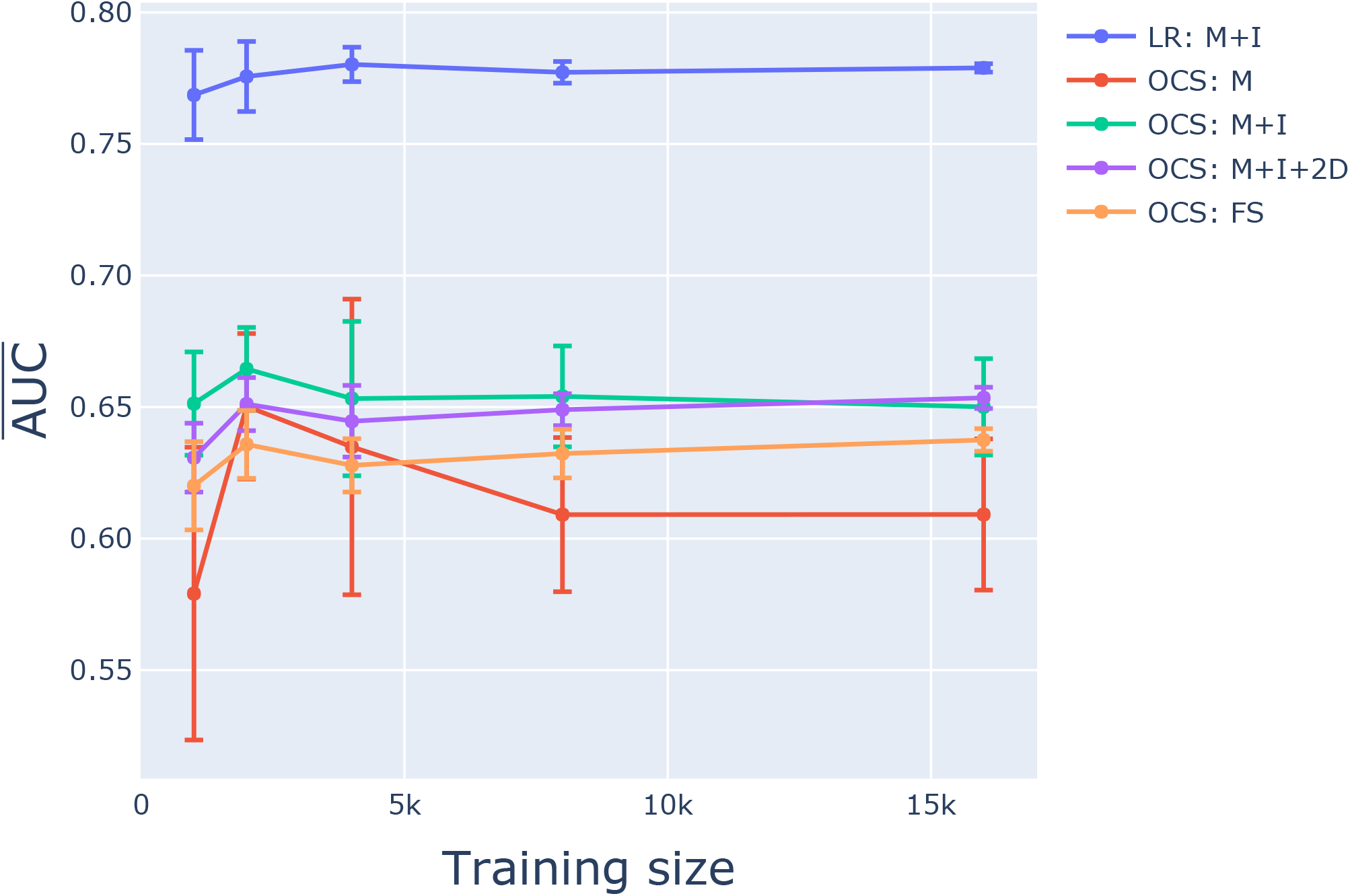
The one-class SVM classifier performed worse than all logistic regression classifiers. x-axis is the size of the training sample, y-axis is the 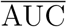 and standard error were calculated as per Fig S2.

**Figure S7:**
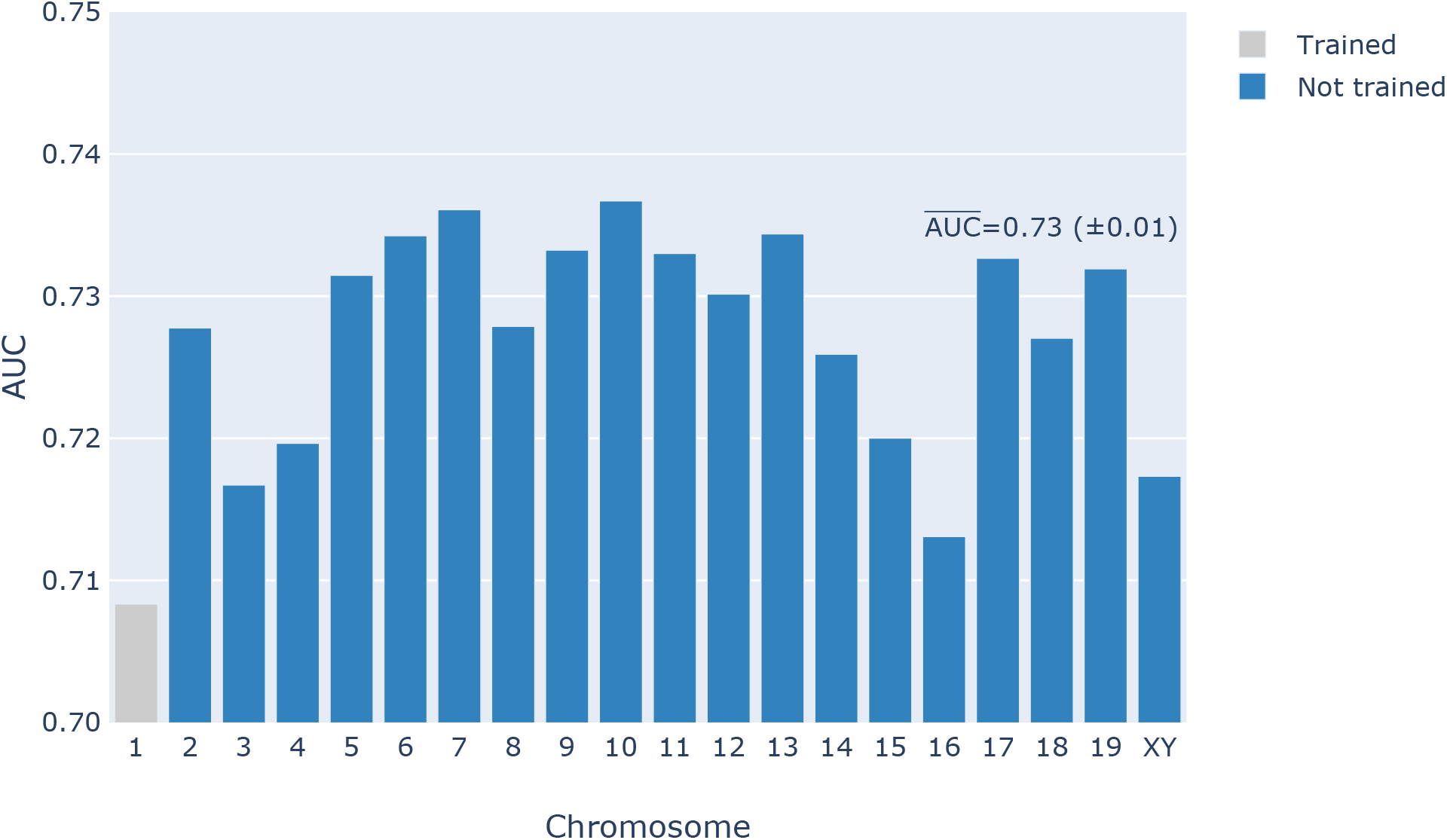
The one-class SVM classifier performed worse than the logistic regression classifier on the entire genome. The classifier was trained on ~1,000 mutations from chromosome 1 using a 5-mer M+I feature set. The 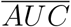 score from the chromosomes not used for training is shown on the figure.

**Figure S8:**
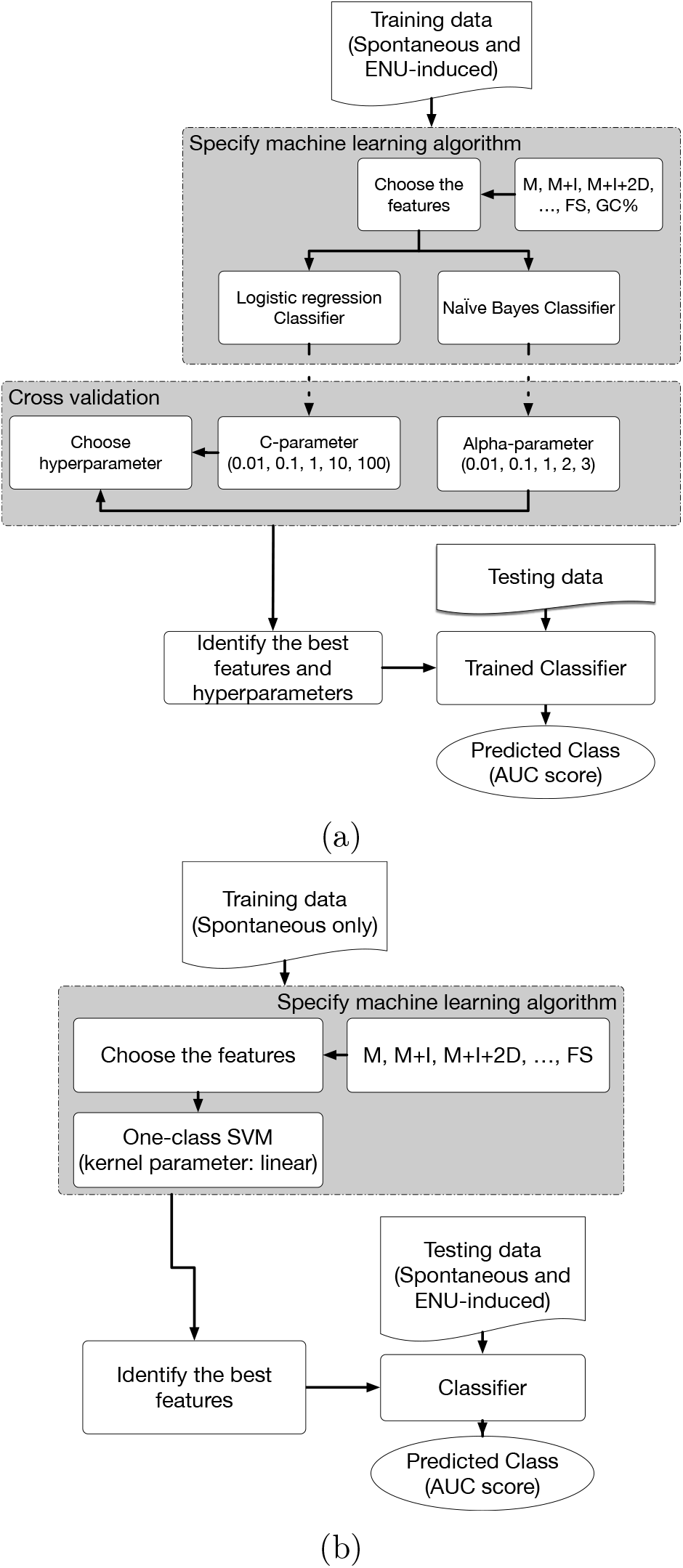
Overview of classifier algorithm evaluation. (a) Two-class classification includes labeled spontaneous and ENU-induced germline point mutations in the training data. (b) One-class classification includes only spontaneous germline point mutations in the training data. For both approaches, training data was limited to mutations occurring on mouse chromosome 1.

**Figure S9:**
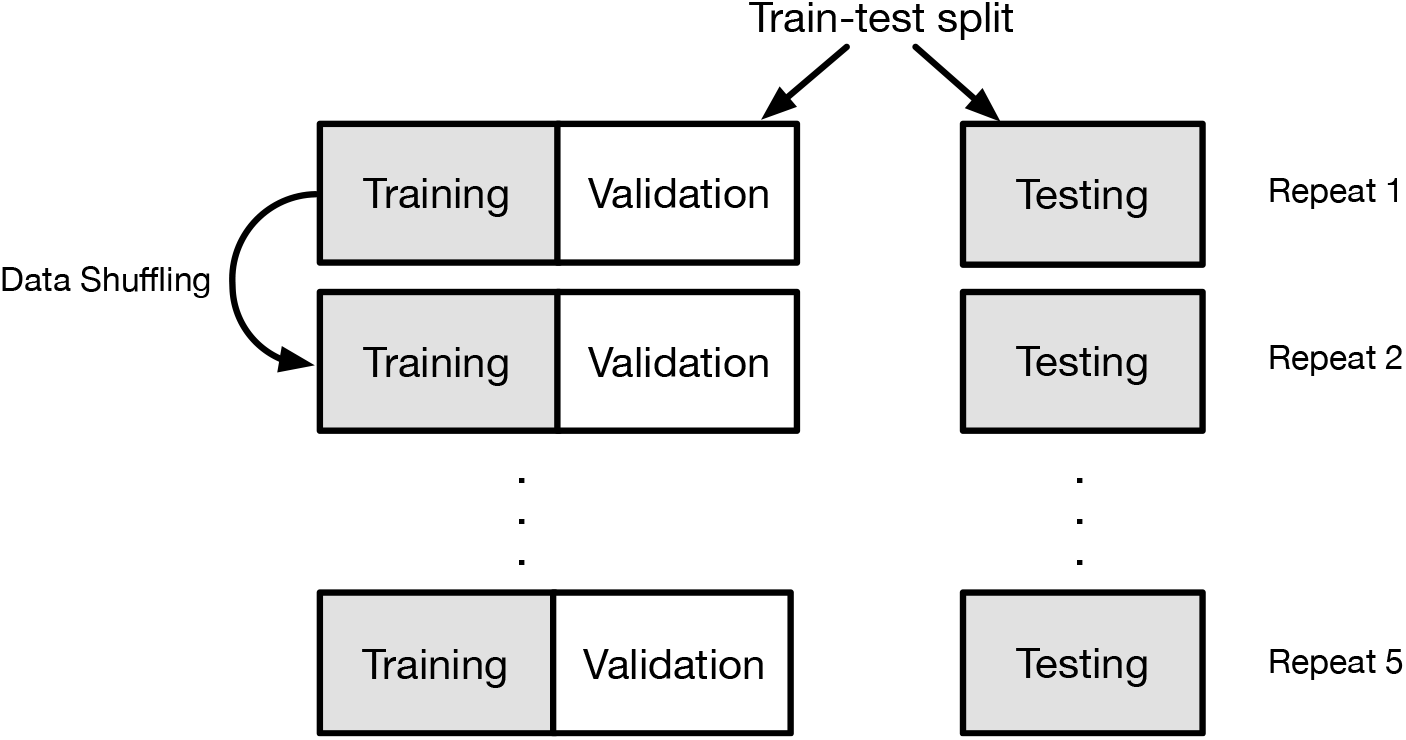
Procedure of cross validation. For each cross validation iteration, the data were shuffled and then divided into three segments, one for training, one for validation and the third one for testing. For each experiment, performance of algorithms with different hyperparameter were compared. The best algorithm for the available data was saved. The process was repeated 10 times.

## References

Aksoy, S. and Haralick, R. M. (2001). Feature normalization and likelihood-based similarity measures for image retrieval. Pattern recognition letters, 22(5):563–582.

Alexandrov, L. B., Nik-Zainal, S., Wedge, D. C., Aparicio, S. A., Behjati, S., Biankin, A. V., Bignell, G. R., Bolli, N., Borg, A., Børresen-Dale, A.-L., et al. (2013). Signatures of mutational processes in human cancer. Nature.

Álvarez, L., Comendador, M., and Sierra, L. (2003). Effect of nucleotide excision repair on ENU-induced mutation in female germ cells of *Drosophila melanogaster*. Environmental and Molecular Mutagenesis, 41(4):270–279.

Andrews, T. D., Whittle, B., Field, M., Balakishnan, B., Zhang, Y., Shao, Y., Cho, V., Kirk, M., Singh, M., Xia, Y., et al. (2012). Massively parallel sequencing of the mouse exome to accurately identify rare, induced mutations: an immediate source for thousands of new mouse models. Open biology, 2(5):120061.

Barbaric, I., Wells, S., Russ, A., and Dear, T. N. (2007). Spectrum of enu-induced mutations in phenotype-driven and gene-driven screens in the mouse. Environmental and molecular mutagenesis, 48(2):124–142.

Bauer, D. C., McMorran, B. J., Foote, S. J., and Burgio, G. (2015). Genome-wide analysis of chemically induced mutations in mouse in phenotype-driven screens. BMC genomics, 16(1):1.

Ben-Hur, A., Ong, C. S., Sonnenburg, S., Schölkopf, B., and Rätsch, G. (2008). Support vector machines and kernels for computational biology. PLoS Comput Biol, 4(10):e1000173.

Bokulich, N. A., Kaehler, B. D., Rideout, J. R., Dillon, M., Bolyen, E., Knight, R., Huttley, G. A., and Gregory Caporaso, J. (2018). Optimizing taxonomic classification of marker-gene amplicon sequences with qiime 2’s q2-feature-classifier plugin. Microbiome, 6(1):90.

Bühlmann, P. and Van De Geer, S. (2011). Statistics for high-dimensional data: methods, theory and applications. Springer Science & Business Media.

Chahwan, R., Edelmann, W., Scharff, M. D., and Roa, S. (2012). AIDing antibody diversity by error-prone mismatch repair. Semin. Immunol., 24(4):293–300.

Chen, T. and Guestrin, C. (2016). XGBoost: A scalable tree boosting system. In Proceedings of the 22nd ACM SIGKDD International Conference on Knowledge Discovery and Data Mining, KDD ’16, pages 785–794, New York, NY USA. ACM.

Davis, J. and Goadrich, M. (2006). The relationship between precision-recall and roc curves. In Proceedings of the 23rd International Conference on Machine Learning.

Hainaut, P. and Pfeifer, G. P. (2001). Patterns of p53 g→t transversions in lung cancers reflect the primary mutagenic signature of dna-damage by tobacco smoke. Carcinogenesis, 22(3):367–374.

Haixiang, G., Yijing, L., Shang, J., Mingyun, G., Yuanyue, H., and Bing, G. (2017). Learning from class-imbalanced data: Review of methods and applications. Expert Systems With Applications, 73(73):220–239.

Hasegawa, M., Kishino, H., and Yano, T.-a. (1985). Dating of the human-ape splitting by a molecular clock of mitochondrial dna. Journal of molecular evolution, 22(2):160–174.

Hellmann, I., Prüfer, K., Ji, H., Zody, M. C., Pääbo, S., and Ptak, S. E. (2005). Why do human diversity levels vary at a megabase scale? Genome research, 15(9):1222–1231.

Hodgkinson, A. and Eyre-Walker, A. (2011). Variation in the mutation rate across mammalian genomes. Nature Reviews Genetics, 12(11):756–766.

Holm, S. (1979). A simple sequentially rejective multiple test procedure. Scandinavian Journal of Statistics, 6(2):65–70.

Huttley, G. A., Jakobsen, I. B., Wilson, S. R., and Easteal, S. (2000). How important is dna replication for mutagenesis? Molecular biology and evolution, 17(6):929–937.

James, G., Witten, D., Hastie, T., and Tibshirani, R. (2013). An introduction to statistical learning, volume 6. Springer.

Justice, M. J., Noveroske, J. K., Weber, J. S., Zheng, B., and Bradley, A. (1999). Mouse enu mutagenesis. Human molecular genetics, 8(10):1955–1963.

King, G. and Zeng, L. (2001). Logistic Regression in Rare Events Data. Political Analysis, 9(02):137–163.

Knight, R., Maxwell, P., Birmingham, A., Carnes, J., Caporaso, J. G., Easton, B. C., Eaton, M., Hamady, M., Lindsay, H., Liu, Z., et al. (2007). Pycogent: a toolkit for making sense from sequence. Genome Biol, 8(8):R171.

Krawczak, M., Ball, E. V., and Cooper, D. N. (1998). Neighboring-nucleotide effects on the rates of germ-line single-base-pair substitution in human genes. The American Journal of Human Genetics, 63(2):474–488.

Lee, J., Cox, B. D., Daly, C. M. S., Lee, C., Nuckels, R. J., Tittle, R. K., Uribe, R. A., and Gross, J. M. (2012). An ENU Mutagenesis Screen in Zebrafish for Visual System Mutants Identifies a Novel Splice-Acceptor Site Mutation in *patched2* that Results in Colobomas. Investigative Opthalmology & Visual Science, 53(13):8214.

Meunier, J. and Duret, L. (2004). Recombination drives the evolution of gc-content in the human genome. Molecular biology and evolution, 21(6):984–990.

Mukherjee, S., Tamayo, P., Rogers, S., Rifkin, R., Engle, A., Campbell, C., Golub, T. R., and Mesirov, J. P. (2003). Estimating dataset size requirements for classifying dna microarray data. Journal of computational biology, 10(2):119–142.

Ng, A. Y. and Jordan, M. I. (2002). On discriminative vs. generative classifiers: A comparison of logistic regression and naive bayes. In Advances in neural information processing systems, pages 841–848.

Noveroske, J., Weber, J., and Justice, M. (2000). The mutagenic action of n-ethyl-n-nitrosourea in the mouse. Mammalian genome, 11(7):478–483.

Peckham, H. E., Thurman, R. E., Fu, Y., Stamatoyannopoulos, J. A., Noble, W. S., Struhl, K., and Weng, Z. (2007). Nucleosome positioning signals in genomic DNA. Genome Research, 17:1170–1177.

Pedregosa, F., Varoquaux, G., Gramfort, A., Michel, V., Thirion, B., Grisel, O., Blondel, M., Prettenhofer, P., Weiss, R., Dubourg, V., Vanderplas, J., Passos, A., Cournapeau, D., Brucher, M., Perrot, M., and Duchesnay, E. (2011). Scikit-learn: Machine learning in Python. Journal of Machine Learning Research, 12:2825–2830.

Pfeifer, G. P., You, Y.-H., and Besaratinia, A. (2005). Mutations induced by ultraviolet light. Mutation Research/Fundamental and Molecular Mechanisms of Mutagenesis, 571(1):19–31.

Pleasance, E. D., Cheetham, R. K., Stephens, P. J., McBride, D. J., Humphray, S. J., Greenman, C. D., Varela, I., Lin, M.-L., Ordóñez, G. R., Bignell, G. R., Ye, K., Alipaz, J., Bauer, M. J., Beare, D., Butler, A., Carter, R. J., Chen, L., Cox, A. J., Edkins, S., Kokko-Gonzales, P. I., Gormley, N. A., Grocock, R. J., Haudenschild, C. D., Hims, M. M., James, T., Jia, M., Kingsbury, Z., Leroy, C., Marshall, J., Menzies, A., Mudie, L. J., Ning, Z., Royce, T., Schulz-Trieglaff, O. B., Spiridou, A., Stebbings, L. A., Szajkowski, L., Teague, J., Williamson, D., Chin, L., Ross, M. T., Campbell, P. J., Bentley, D. R., Futreal, P. A., and Stratton, M. R. (2010). A comprehensive catalogue of somatic mutations from a human cancer genome. Nature, 463(7278):191–196.

Prosperi, M. C., Altmann, A., Rosen-Zvi, M., Aharoni, E., Borgulya, G., Bazso, F., Sönnerborg, A., Schülter, E., Struck, D., Ulivi, G., et al. (2009). Investigation of expert rule bases, logistic regression, and non-linear machine learning techniques for predicting response to antiretroviral treatment. Antivir Ther, 14(3):433–42.

Shiraishi, Y., Tremmel, G., Miyano, S., and Stephens, M. (2015). A simple model-based approach to inferring and visualizing cancer mutation signatures. PLoS Genet, 11(12):e1005657.

Shrivastav, N., Li, D., and Essigmann, J. M. (2010). Chemical biology of mutagenesis and dna repair: cellular responses to dna alkylation. Carcinogenesis, 31(1):59–70.

Sonnenburg, S. (2008). Machine Learning for Genomic Sequence Analysis-Dissertation. PhD thesis, Berlin Institute of Technology.

Stottmann, R. and Beier, D. (2014). ENU Mutagenesis in the Mouse. Current protocols in human genetics, 82:15.4.1–10.

Svejstrup, J. Q. (2002). Mechanisms of transcription-coupled DNA repair. Nat Rev Mol Cell Biol, 3(1):21–29.

Takahasi, K. R., Sakuraba, Y., and Gondo, Y. (2007). Mutational pattern and frequency of induced nucleotide changes in mouse enu mutagenesis. BMC molecular biology, 8(1):1.

Viel, A., Bruselles, A., Meccia, E., Fornasarig, M., Quaia, M., Canzonieri, V., Policicchio, E., Urso, E. D., Agostini, M., Genuardi, M., et al. (2017). A specific mutational signature associated with dna 8-oxoguanine persistence in mutyh-defective colorectal cancer. EBioMedicine.

Wålinder, A. (2014). Evaluation of logistic regression and random forest classification based on prediction accuracy and metadata analysis.

Yakovchuk, P., Protozanova, E., and Frank-Kamenetskii, M. D. (2006). Base-stacking and basepairing contributions into thermal stability of the DNA double helix. Nucleic Acids Research, 34(2):564–574.

Yang, Z., Kumar, S., and Nei, M. (1995). A new method of inference of ancestral nucleotide and amino acid sequences. Genetics, 141(4):1641–1650.

Zhu, Y., Neeman, T., Yap, V. B., and Huttley, G. A. (2017). Statistical methods for identifying sequence motifs affecting point mutations. Genetics, 205(2):843–856.

