## Supplementary Tables and Figures for "Machine learning techniques for classifying the mutagenic origins of point mutations"

October 31, 2019

| **Direction** | **Class** | **RET** |
|---|---|---|
| T→C | ENU | -0.047 |
| A→T | Spontaneous | -0.036 |
| G→T | Spontaneous | -0.036 |
| T→A | Spontaneous | -0.035 |
| A→G | ENU | -0.035 |
| C→A | Spontaneous | -0.034 |
| G→A | ENU | -0.025 |
| C→T | ENU | -0.021 |
| A→C | ENU | -0.018 |
| T→G | ENU | -0.007 |
| G→C | ENU | -0.001 |
| C→G | ENU | -0.001 |
| T→G | Spontaneous | 0.009 |
| C→G | Spontaneous | 0.022 |
| C→T | Spontaneous | 0.022 |
| G→C | Spontaneous | 0.023 |
| G→A | Spontaneous | 0.027 |
| A→C | Spontaneous | 0.027 |
| A→G | Spontaneous | 0.039 |
| C→A | ENU | 0.052 |
| T→C | Spontaneous | 0.055 |
| G→T | ENU | 0.063 |
| T→A | ENU | 0.066 |
| A→T | ENU | 0.067 |

Table S1: Comparison of mutation spectra between Spontaneous and ENU-induced germline point mutations. RET values are proportional to deviance generated from the log-linear model (Zhu et al., 2017), and $p$-value are obtained from the $\chi^2$ distribution. All $p$-values were below the limit of detection.

| Mutation direction | 1st-order | 2nd-order | 3rd-order | 4th-order |
|---|---|---|---|---|
| A→C | 4 | 5 | 3 | 0 |
| A→G | 4 | 5 | 4 | 1 |
| A→T | 4 | 5 | 2 | 1 |
| C→A | 4 | 6 | 4 | 1 |
| C→T | 4 | 5 | 4 | 1 |
| G→A | 4 | 5 | 4 | 1 |
| G→T | 4 | 5 | 2 | 1 |
| T→A | 4 | 6 | 2 | 0 |
| T→C | 4 | 6 | 4 | 0 |
| T→G | 4 | 5 | 3 | 1 |

Table S2: Number of positions showing significant differences between ENU-induced and spontaneous germline point mutations from analysis of 5-mers. A $p$-value $\leq$ 0.05 was classified as significant. $p$-values were from the log-linear analysis.

| **Direction** | **RE**$_{max}$(1) | **RE Dist.** | **p-val Dist.** |
|---|---|---|---|
| A→C | 0.0374 | 6 | 10 |
| A→G | 0.0402 | 4 | 10 |
| A→T | 0.0638 | 2 | 10 |
| C→A | 0.0632 | 2 | 10 |
| C→T | 0.0703 | 2 | 10 |
| G→A | 0.0710 | 2 | 10 |
| G→T | 0.0624 | 2 | 10 |
| T→A | 0.0606 | 2 | 10 |
| T→C | 0.0395 | 4 | 10 |
| T→G | 0.0373 | 6 | 10 |

(a) ENU-induced

| **Direction** | **RE**$_{max}$(1) | **RE Dist.** | **p-val Dist.** |
|---|---|---|---|
| A→C | 0.0047 | 8 | 10 |
| A→G | 0.0118 | 3 | 10 |
| A→T | 0.0194 | 3 | 10 |
| C→A | 0.0332 | 4 | 10 |
| C→T | 0.0505 | 1 | 10 |
| G→A | 0.0508 | 1 | 10 |
| G→T | 0.0351 | 3 | 10 |
| T→A | 0.0117 | 2 | 10 |
| T→C | 0.0152 | 2 | 10 |
| T→G | 0.0148 | 2 | 10 |

(b) Spontaneous

Table S3: Longer range neighbourhood effect log-linear analyses results of (a) ENU-induced mutations and (b) germline spontaneous mutations. For both subtables, the most distant positions from the mutation with RE(1) $\geq$ 10% of RE$_{max}$(1). RE(1) is the first order RE for the position, and RE$_{max}$(1) the largest RE from a first order effect for the surveyed positions. RE Dist. is the furthest position with an RE value $\geq 0.1 imes$RE$_{max}$. p-val Dist. is the corresponding distance based on the $p$-value$\leq$ 0.05. As the analysis was limited to a flank size of 10bp either side of the mutating base, the maximum possible distance is 10.

| **Chromosome** | **ENU-induced** | **Spontaneous** |
|---|---|---|
| 1 | 16,977 | 17,848 |
| 2 | 21,100 | 20,051 |
| 3 | 11,228 | 11,713 |
| 4 | 13,973 | 16,936 |
| 5 | 14,509 | 16,028 |
| 6 | 13,039 | 12,097 |
| 7 | 20,864 | 19,161 |
| 8 | 11,232 | 13,465 |
| 9 | 14,010 | 15,662 |
| 10 | 11,315 | 12,641 |
| 11 | 17,101 | 19,626 |
| 12 | 8,022 | 8,817 |
| 13 | 9,085 | 8,939 |
| 14 | 8,395 | 8,868 |
| 15 | 9,342 | 11,079 |
| 16 | 7,266 | 8,117 |
| 17 | 11,981 | 12,168 |
| 18 | 6,356 | 7,732 |
| 19 | 7,529 | 8,635 |
| XY | 853 | 5,097 |

Table S4: By-chromosome sample sizes of genetic variants from the ENU induced and spontaneous germline mutations.

| **Feature set** | **Training size** | **mean(auc)** | **std(auc)** | **min(auc)** | **max(auc)** |
|---:|---:|:---:|:---:|:---:|:---:|
| FS | 1,000 | 0.759 | 0.019 | 0.723 | 0.788 |
| FS | 2,000 | 0.775 | 0.015 | 0.745 | 0.800 |
| FS | 4,000 | 0.782 | 0.007 | 0.771 | 0.790 |
| FS | 8,000 | 0.782 | 0.004 | 0.776 | 0.790 |
| FS | 16,000 | 0.788 | 0.002 | 0.784 | 0.792 |
| M+I | 1,000 | 0.769 | 0.017 | 0.746 | 0.793 |
| M+I | 2,000 | 0.776 | 0.013 | 0.752 | 0.799 |
| M+I | 4,000 | 0.780 | 0.007 | 0.771 | 0.787 |
| M+I | 8,000 | 0.777 | 0.004 | 0.771 | 0.783 |
| M+I | 16,000 | 0.779 | 0.002 | 0.776 | 0.782 |
| M+I+2D | 1,000 | 0.757 | 0.022 | 0.723 | 0.792 |
| M+I+2D | 2,000 | 0.774 | 0.015 | 0.747 | 0.800 |
| M+I+2D | 4,000 | 0.782 | 0.005 | 0.772 | 0.788 |
| M+I+2D | 8,000 | 0.783 | 0.005 | 0.777 | 0.791 |
| M+I+2D | 16,000 | 0.786 | 0.002 | 0.781 | 0.789 |
| M+I+2Dp | 1,000 | 0.757 | 0.019 | 0.734 | 0.787 |
| M+I+2Dp | 2,000 | 0.773 | 0.015 | 0.746 | 0.802 |
| M+I+2Dp | 4,000 | 0.782 | 0.006 | 0.773 | 0.788 |
| M+I+2Dp | 8,000 | 0.782 | 0.004 | 0.777 | 0.790 |
| M+I+2Dp | 16,000 | 0.784 | 0.002 | 0.780 | 0.787 |

Table S5: Summary of AUC scores from LR classifiers using 7-mers.

| **Feature set** | **Training size** | **mean(auc)** | **std(auc)** | **min(auc)** | **max(auc)** |
|---:|---:|:---:|:---:|:---:|:---:|
| FS | 1,000 | 0.760 | 0.017 | 0.733 | 0.787 |
| FS | 2,000 | 0.768 | 0.014 | 0.743 | 0.795 |
| FS | 4,000 | 0.774 | 0.008 | 0.762 | 0.785 |
| FS | 8,000 | 0.771 | 0.004 | 0.765 | 0.777 |
| FS | 16,000 | 0.773 | 0.002 | 0.769 | 0.775 |
| M+I | 1,000 | 0.765 | 0.016 | 0.738 | 0.788 |
| M+I | 2,000 | 0.769 | 0.013 | 0.745 | 0.792 |
| M+I | 4,000 | 0.773 | 0.008 | 0.763 | 0.785 |
| M+I | 8,000 | 0.769 | 0.003 | 0.765 | 0.774 |
| M+I | 16,000 | 0.771 | 0.002 | 0.768 | 0.774 |

Table S6: Summary of AUC scores from LR classifiers using 3-mers.

| Feature set | Training size | mean(auc) | std(auc) | min(auc) | max(auc) |
|---|---|---|---|---|---|
| FS | 1,000 | 0.752 | 0.017 | 0.727 | 0.775 |
| FS | 2,000 | 0.770 | 0.014 | 0.742 | 0.793 |
| FS | 4,000 | 0.778 | 0.008 | 0.767 | 0.787 |
| FS | 8,000 | 0.777 | 0.004 | 0.772 | 0.786 |
| FS | 16,000 | 0.779 | 0.002 | 0.777 | 0.782 |
| M+I | 1,000 | 0.763 | 0.018 | 0.741 | 0.788 |
| M+I | 2,000 | 0.771 | 0.014 | 0.745 | 0.795 |
| M+I | 4,000 | 0.776 | 0.008 | 0.763 | 0.787 |
| M+I | 8,000 | 0.772 | 0.004 | 0.767 | 0.778 |
| M+I | 16,000 | 0.774 | 0.002 | 0.772 | 0.777 |
| M+I+2D | 1,000 | 0.753 | 0.023 | 0.717 | 0.789 |
| M+I+2D | 2,000 | 0.769 | 0.014 | 0.742 | 0.793 |
| M+I+2D | 4,000 | 0.778 | 0.008 | 0.766 | 0.786 |
| M+I+2D | 8,000 | 0.777 | 0.004 | 0.772 | 0.786 |
| M+I+2D | 16,000 | 0.779 | 0.001 | 0.776 | 0.781 |
| M+I+2Dp | 1,000 | 0.758 | 0.019 | 0.728 | 0.784 |
| M+I+2Dp | 2,000 | 0.770 | 0.015 | 0.742 | 0.797 |
| M+I+2Dp | 4,000 | 0.778 | 0.007 | 0.767 | 0.785 |
| M+I+2Dp | 8,000 | 0.776 | 0.004 | 0.772 | 0.786 |
| M+I+2Dp | 16,000 | 0.778 | 0.002 | 0.775 | 0.780 |

Table S7: Summary of AUC scores from LR classifiers using 5-mers.

| Feature set | Training size | mean(auc) | std(auc) | min(auc) | max(auc) |
|---|---|---|---|---|---|
| M+I | 1,000 | 0.765 | 0.017 | 0.730 | 0.792 |
| M+I | 2,000 | 0.791 | 0.013 | 0.763 | 0.804 |
| M+I | 4,000 | 0.804 | 0.006 | 0.796 | 0.813 |
| M+I | 8,000 | 0.810 | 0.003 | 0.807 | 0.817 |
| M+I | 16,000 | 0.814 | 0.002 | 0.812 | 0.817 |
| M+I+2Dp | 1,000 | 0.763 | 0.016 | 0.737 | 0.783 |
| M+I+2Dp | 2,000 | 0.790 | 0.012 | 0.765 | 0.806 |
| M+I+2Dp | 4,000 | 0.808 | 0.005 | 0.798 | 0.813 |
| M+I+2Dp | 8,000 | 0.820 | 0.004 | 0.814 | 0.825 |
| M+I+2Dp | 16,000 | 0.827 | 0.002 | 0.825 | 0.831 |

Table S8: Summary of AUC scores from LR classifiers using 59-mers.

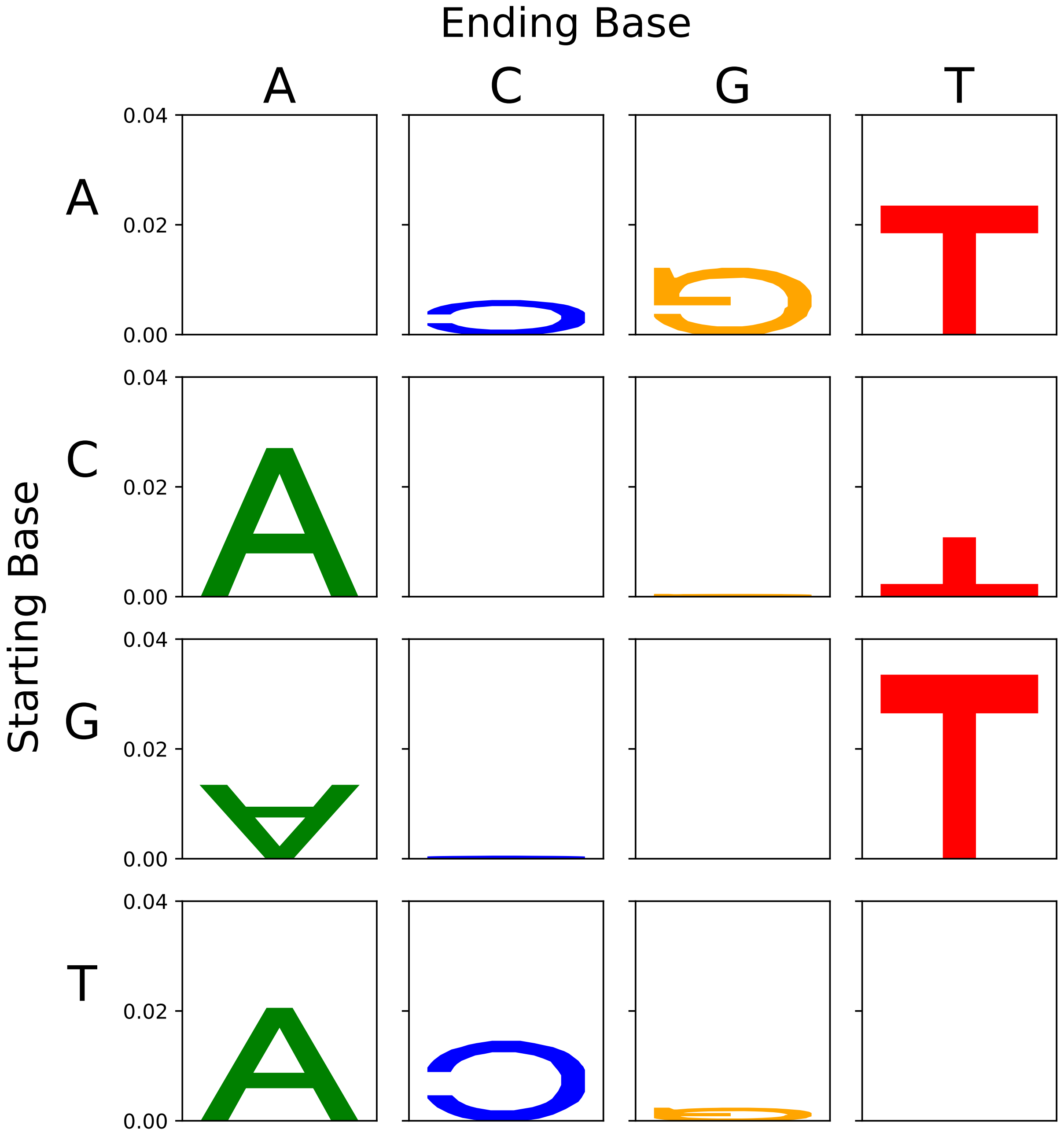


Figure S1: Confirmation of the mutation spectra difference between the ENU-induced and spontaneous germline mutations. Starting and Ending Base correspond to X, Y respectively in X→Y. The y-axis is RE from the spectra hypothesis test and letter heights are as for the mutation motif logo. Letters in the normal orientation indicate an excess of that mutation direction in ENU-induced mutations relative to the spontaneous mutations. Inverted letters indicate a deficit in ENU-induced mutations relative to the spontaneous mutations. See Zhu et al. (2017) for a more detailed description of the log-linear models.

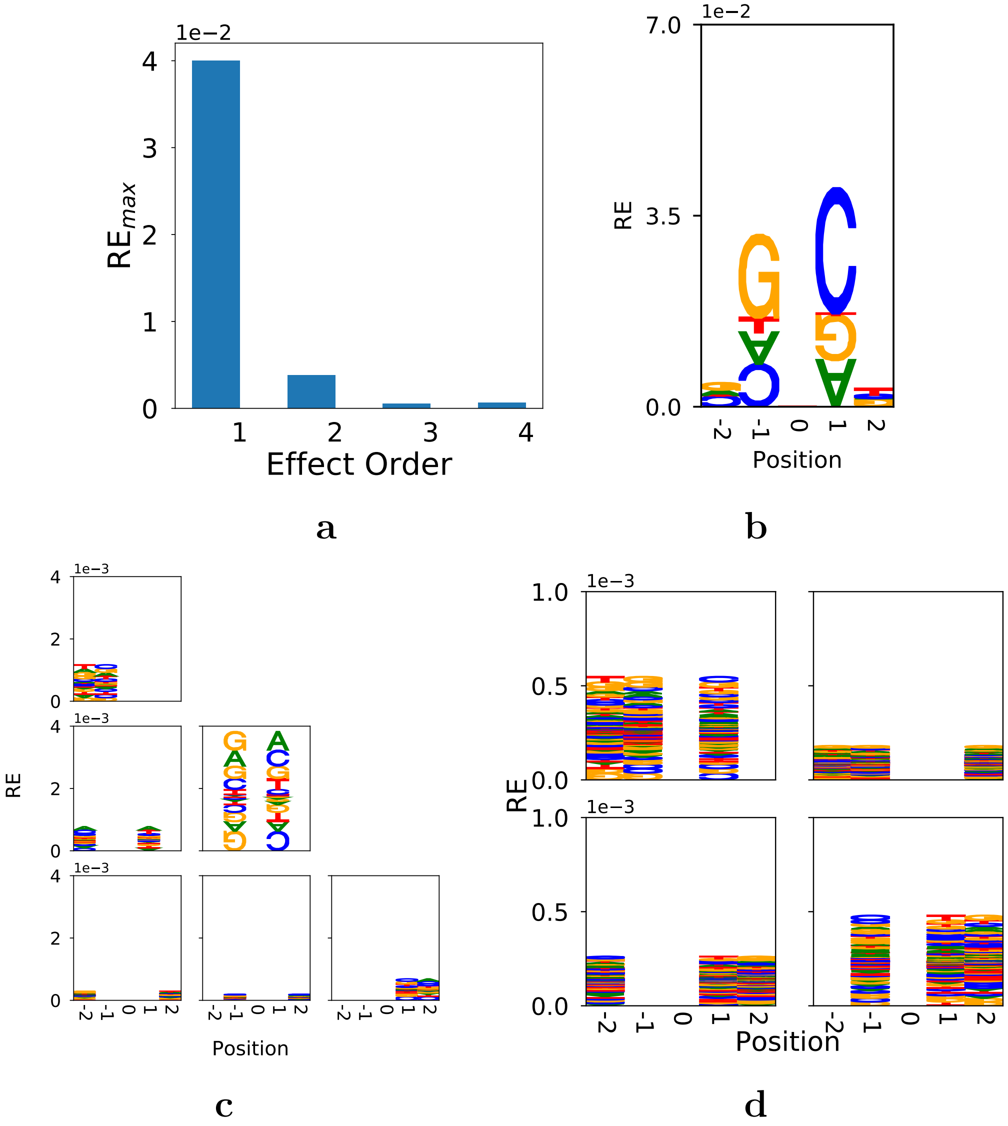


Figure S2: Independent and second-order position effects dominate ENU-induced A→G point mutations. Note also that RE is largest for dependent effects among positions that are positions physically contiguous and overlap the mutated position at index 0. (a) Summary of the strength of associations by effect order. $RE_{max}$ is the maximum RE from any analysis for the indicated order. (b) The independent, or first-order, effects. (c) Second-order effects. (d) third-order effects.

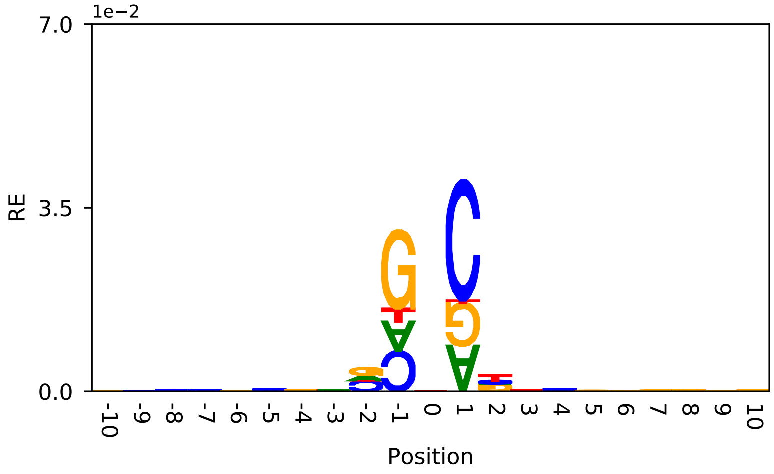


(a) ENU-induced

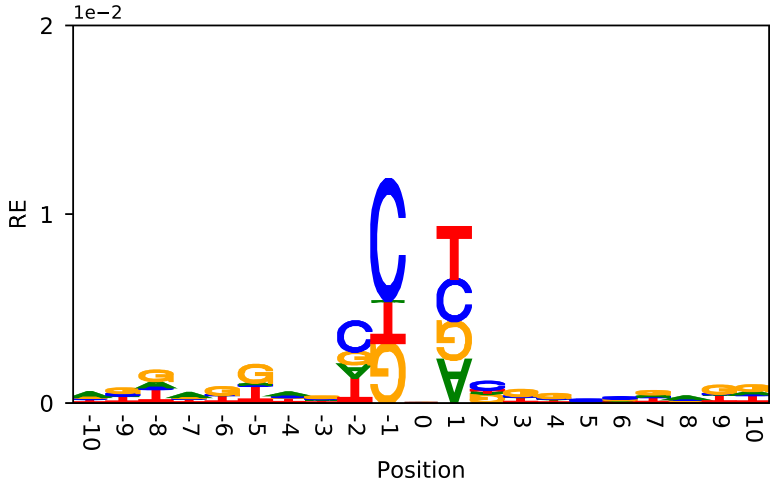


(b) Spontaneous

Figure S3: The physical extent of neighbourhood effects in the mouse. Mutation motifs are drawn from the results of the log-linear analysis of first-order effects (summarised in Table S3). (a) ENU-induced germline mutations and (b) Spontaneous germline mutations.

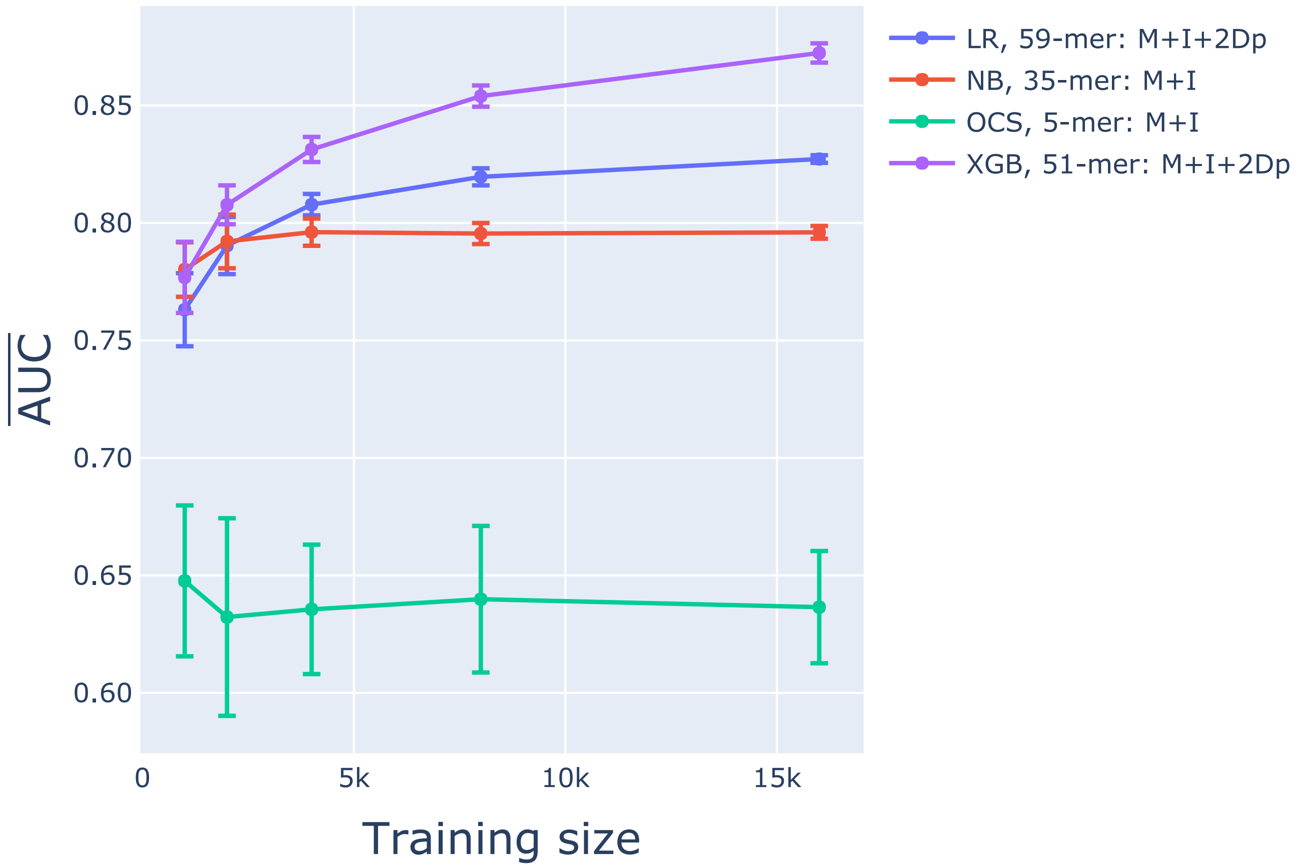


Figure S4: Comparison of the top-ranked classifiers. For a given algorithm, the classifier set with the largest AUC from an individual replicate was chosen as the best classifier. x-axis is the size of the training sample, y-axis is the mean AUC and error bars were calculated from the 10 chromosome 1 training samples. The algorithm, $k$-mer and feature set were as indicated.

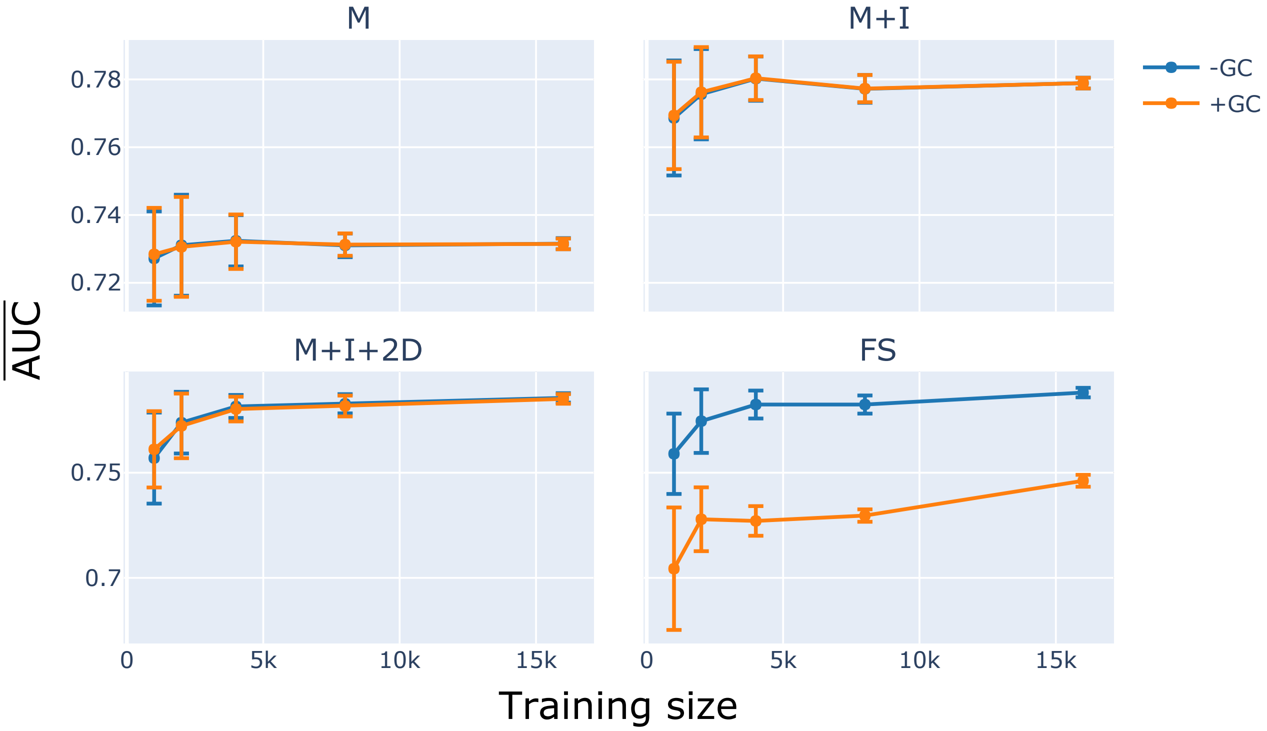


Figure S5: Inclusion of GC% did not improve performance when categorical neighbourhood features were included.

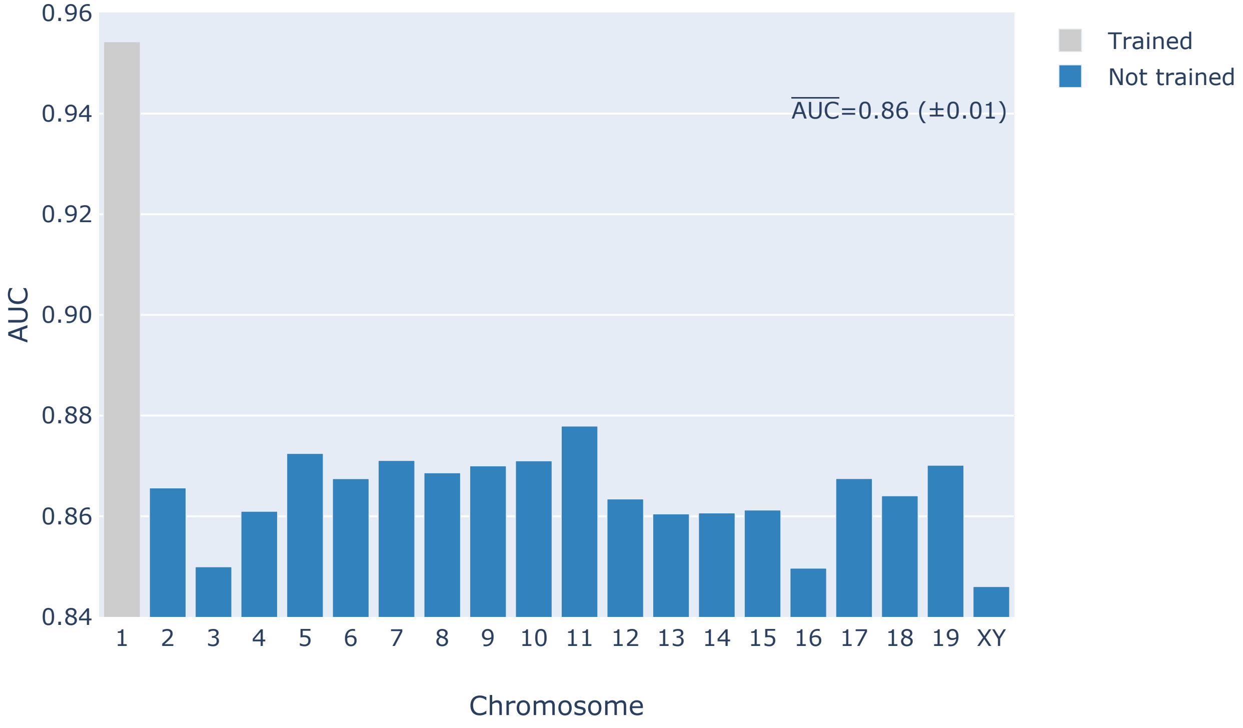


Figure S6: Per chromosome classification performance on the mouse genome of the best XGB classifier. The classifier was trained on 16,000 mutations from chromosome 1 using a 51-mer M+I+2Dp feature set. The $\overline{AUC}$ score from the chromosomes not used for training is shown on the figure.

### References

Zhu, Y., Neeman, T., Yap, V. B., and Huttley, G. A. (2017). Statistical methods for identifying sequence motifs affecting point mutations. *Genetics*, 205(2):843–856.
